# Towards an evolutionary baseline model of *Plasmodium falciparum* for population-genomic inference

**DOI:** 10.64898/2025.12.20.695730

**Authors:** Cobi M. Henry, Jacob I. Marsh, Austin Daigle, James Crescenzi, Jessica T. Lin, Jeffrey A. Bailey, Parul Johri

## Abstract

Malaria has caused over 15.7 million deaths in the 21^st^ century and was responsible for ∼600 thousand deaths globally in 2023 alone. Although many effective antimalarial drugs have been developed and widely adopted to reduce the occurrence and severity of the disease, recurrent resistance to the frontline treatment has been of major concern. Multiple drug resistance alleles at intermediate and high allele frequency have been identified in specific Asian and African populations of *P. falciparum*, the deadliest malaria parasite. With the improvement in throughput of sequencing technologies and global efforts such as the MalariaGEN project to build genomic surveillance, we now have access to tens of thousands of genomes of *P. falciparum* from across the world. With this data, it is becoming increasingly possible to employ powerful population genetics approaches to understand the selective pressures and demographic history of the parasite. While several empirically motivated outlier-based approaches have been employed to identify targets of drug resistance, there is a lack of a framework that jointly accounts for the multiple concurrent processes occurring in natural populations of *P. falciparum*. We argue that a baseline evolutionary model that accounts for simultaneously acting evolutionary processes is needed to understand patterns of genomic variation in *P. falciparum* populations. Here, we identify key components essential for building such a baseline model for the malaria-causing pathogen. The development of an appropriate null model will be important to test evolutionary hypotheses using genomic datasets, will provide a path forward to improve the accuracy of inference of evolutionary parameters, and will help identify new gene candidates involved in drug resistance.

## INTRODUCTION

*Plasmodium falciparum*, a unicellular protozoan parasite that is responsible for malaria, accounts for over 90% of malaria mortality worldwide (Snow 2015; World Health Organization 2024). It causes the most severe and fatal outcomes among all malaria-causing species, with approximately half of the world’s population at risk of contracting the disease (Centers for Disease Control and Prevention 2024). In the 20th century alone, *P. falciparum* claimed an estimated 150-300 million lives and continues to kill approximately half a million people per year (Arrow et al. 2004; World Health Organization 2024). Despite continuous efforts to develop effective antimalarial drugs, *P. falciparum* has repeatedly developed wide-spread resistance to frontline treatments, including chloroquine in the 1960s and sulfadoxine–pyrimethamine in the 1980s (Payne 1987; Basu et al. 2025). Most recently, acquired partial resistance to current frontline artemisinin-based combination therapy (ACT) has emerged, jeopardizing recent advancements in malaria control (Phyo et al. 2016; Rosenthal et al. 2024). Given the severity and widespread impact of *P. falciparum* transmission, understanding the evolutionary dynamics shaping its populations has become increasingly critical. With the advent of a large amount of publicly available whole-genome population-genetic data from more than 30,000 genomes of *P.* falciparum (Ahouidi et al. 2021; Abdel Hamid et al. 2023; Abdel Hamid et al. 2025), ∼2000 genomes of *P. vivax* individuals (Adam et al. 2022), and initial effort for *P. malariae* (Popkin-Hall et al. 2024) and *P. ovale* (Carey-Ewend et al. 2024), population-genomic inference is likely to become a widespread approach to understand the evolution of drug resistance, adaptation to the host, and key demographic events in *Plasmodium* (Volkman et al. 2007; Chang et al. 2012; Parobek et al. 2016; Guo et al. 2024).

When a drug-resistance or any other beneficial allele rapidly increases in frequency and/or reaches fixation in the population, it results in molecular signatures at sites linked to the beneficial fixation. Specifically, there is a reduction in genetic diversity, a skew in the site frequency spectrum (SFS), and a unique spatial pattern of linkage disequilibrium (LD) generated at linked sites, also known as selective sweeps (Stephan 2019). Thus, statistical summaries of DNA sequence variation are commonly used to detect candidates of recent fixation of beneficial mutations from population-genetic data (Nielsen 2005). For instance, outlier-based approaches use loci present in the tails of the distribution of summary statistics, like measures of genetic diversity, population differentiation, skew in the SFS relative to expectations under neutrality (Tajima 1989) and haplotype diversity *(e.g.,* Garud et al. 2015), to identify possible gene candidates that have recently experienced sweeps. However, variability in such statistics across the genome is generated not only by a rapid increase in frequency of beneficial mutations or those under balancing selection, but also due to other non-adaptive processes like variation in mutation or recombination rates and simply due to strong bottlenecks, *i.e*., the sharp reduction in the size of a population (Teshima et al. 2006).

There have been two opposing schools of thought when accounting for this issue. One school of thought recommends identifying putative candidates of positive selection by simply selecting all loci present in the tails (1% or 5%) of the distributions of summary statistics (*e.g.*, Carlson et al. 2005; Kelley et al. 2006). This approach has many caveats (Thornton and Jensen 2007), with the two most important ones being that it always yields potential gene candidates even in the absence of any recent selective sweeps (because a fixed fraction of all genes, such as 1% or 5% will always be identified as candidates). Secondly, this approach assumes that loci that have undergone sweeps will be enriched in the tails of the distributions, which is not necessarily true (as shown by Teshima et al. 2006), and thus the false positive rate is unknown. For instance, severe population bottlenecks can drastically increase the false positive rate when detecting selective sweeps using outliers (Crisci et al. 2013), and the effects of purifying selection and recombination rate variation can result in falsely identifying outlier loci even without any beneficial fixations (Johri, Stephan, et al. 2022).

To address these caveats, the second school of thought suggests a model-based approach, wherein putative sweep candidates are identified by first generating an expectation of the distribution of the statistic of interest in the absence of positive selection by explicitly modeling the history of the population of interest (Teshima et al. 2006; Johri, Aquadro, et al. 2022). The caveat of the model-based approach is that model violations or poor model fit may lead to incorrect inferences. We therefore stress the importance of working towards building accurate and realistic models so that we can detect loci under positive or balancing selection with higher sensitivity and specificity so that a fuller picture of the genes responsible for adaptation can be obtained (Johri, Aquadro, et al. 2022; Soni et al. 2023; Soni and Jensen 2024). In fact, while a number of outlier-based approaches have been employed in *Plasmodium* species to identify targets of drug resistance (*e.g.,* Volkman et al. 2007; Parobek et al. 2016), there is a lack of an evolutionary model that jointly accounts for multiple constantly operating evolutionary processes in natural parasite populations, shaping molecular variation across their genomes.

Accounting for an evolutionary baseline model in unicellular pathogenic species can be uniquely challenging for multiple reasons. (a) Unlike human genomes, which are sparsely populated with functionally important genomic elements, genomes of unicellular species are often characterized by highly streamlined genomes (Lynch and Conery 2003), that are gene-rich, making the effects of selection pervasive across the genome. (b) Unicellular species often exhibit some form of clonal reproduction or self-fertilization (Weedall and Hall 2015), which reduces the effectiveness of recombination in their population, in turn increasing the effects of selection across the genome. (c) Finally, pathogenic organisms experience a highly complex demographic history due to continual bottlenecks during host invasions and strong bouts of selection due to exposure to antimicrobial agents (Renzette et al. 2013). Thus, in order to model molecular variation in natural populations of pathogenic species, it becomes especially important to jointly account for the effects of multiple evolutionary factors, like their mating system, skewed distribution of offspring, direct selection, the effects of selection on putatively neutral sites, and complex demography.

We argue that evolutionary baseline models of *Plasmodium falciparum* must account for such factors, in particular, recurrent bottlenecks, realistic genome architecture (density of coding and intergenic regions), selection against deleterious mutations, along with heterogeneity in mutation and recombination rates across the genome Johri, Aquadro, et al. 2022). Such baseline models can then be employed to generate realistic expectations of genomic variation and thus to better identify putative sweep candidates (detailed in Box 4 of Johri, Aquadro, et al. 2022). We here elucidate how such an evolutionary baseline model in *P. falciparum* may be constructed and which population-genetic considerations likely play important roles in evolutionary inference.

With the growing amount of publicly available whole-genome population-genetic data of *P.* falciparum (Ahouidi et al. 2021; Abdel Hamid et al. 2023; Abdel Hamid et al. 2025), there will be many opportunities for performing population-genomic inference. It is therefore imperative that we construct baseline models, which would aid in biologic inference correctly interpreting evolutionary forces shaping genomic variation in human pathogens.

### INFECTION CYCLE

The *P. falciparum* life cycle consists of two main phases: the human phase and the mosquito phase. In the salivary gland of the female mosquito, ∼10 (range: 10^3^-10^5^) haploid sporozoites, or the spore-like, motile phase of the parasite, remain in the salivary gland until the next blood meal (Kappe et al. 2010; Graumans et al. 2020; Andolina et al. 2024; Kanatani et al. 2024). The human phase starts when an infectious female *Anopheles* mosquito bites a human, sporozoites are injected with the saliva into the dermis of the human host (reviewed in Ejigiri and Sinnis 2009). Only ∼20% of sporozoites reach the salivary glands of mosquitoes, and 10^2^-10^3^ are transmitted during the blood meal to the dermis of the human host (Graumans et al. 2020; Andolina et al. 2024; Kanatani et al. 2024). The sporozoites remain motile in the dermis and follow a random path until a portion of them encounters a blood vessel and circulate into the liver. Those that fail to reach the bloodstream are either destroyed in the skin or drained by the lymphatic system (Amino et al. 2006; Yamauchi et al. 2007). On average, the majority of sporozoites that reach the liver take between 1 and 3 hours to leave the skin (Yamauchi et al. 2007). After reaching the liver, during the **hepatocytic (liver cell) stage** (see Figure 1), the surviving sporozoites (∼10 in number, range: 1-10^2^) invade hepatocytes and begin their first round of mitotic asexual reproduction (Mota et al. 2001; Kappe et al. 2010; Vaughan and Kappe 2017; Graumans et al. 2020; Venugopal et al. 2020). After 13-14 rounds of mitosis, the result is a multi-nucleated schizont (cell with tens of thousands of nuclei) containing thousands of haploid daughter cells called merozoites. After 6-7 days of multiplying in the liver, the schizont bursts and releases up to 90,000 (range: 10^4^ – 10^5^) merozoites into the bloodstream (Kappe et al. 2010; Vaughan and Kappe 2017; Graumans et al. 2020), where each of the merozoites rapidly invades a circulating red blood cell (Venugopal et al. 2020).

**Figure 1:**
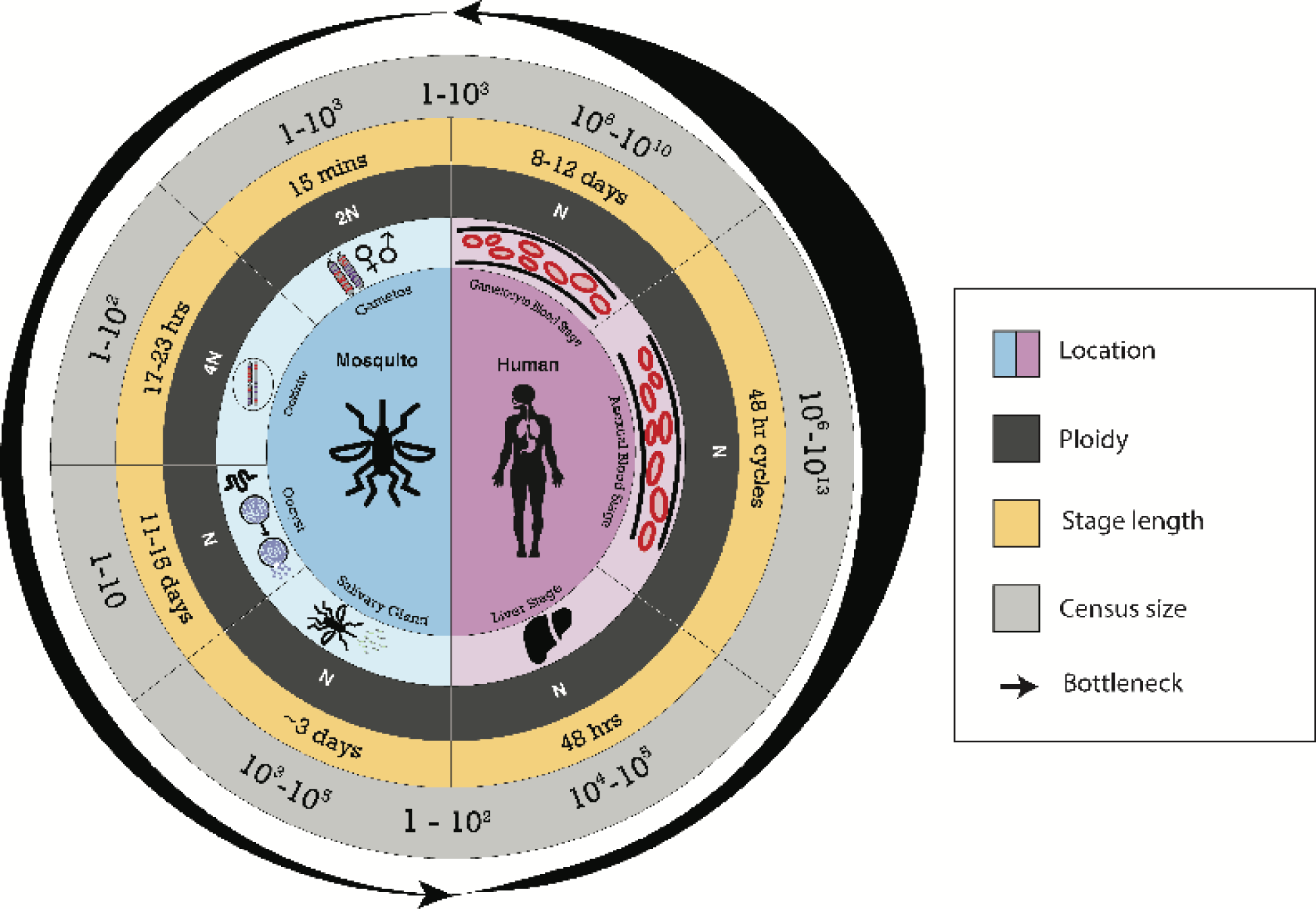
The lifecycle of *Plasmodium falciparum*. The progression of various life stages is displayed in human (purple) and mosquito (blue) hosts. The inner-most circle shows the stage of the lifecycle and the corresponding ploidy of *P. falciparum* in grey circles. Note that the gametocyte blood stage and the asexual blood stage may occur concurrently within a human host as parasites can be in different stages at any given time. The yellow ring displays the approximate time spent in the corresponding stage (not drawn to scale). The outermost circle displays recurrent bottlenecks experienced by *P. falciparum* during the life cycle and the numbers indicate present estimates of the number of cells. The width of the black arrows is correlated to the estimated within-host population sizes.

Within the red blood cells (RBCs), the parasites undergo additional rounds of asexual reproduction over the course of 48-hour cycles (Hawking et al. 1968; Smith et al. 2020), referred to as the **asexual erythrocyte (RBC) stage** (Figure 1), where they multiply into a schizont containing 32 daughter merozoites after 5 rounds of mitosis (Venugopal et al. 2020). At this point, the schizont bursts and these merozoites rapidly invade new RBCs to initiate another replication cycle (Cowman and Crabb 2006). This exponential process allows the parasite population to increaseby approximately 8 to 10-fold every 48 hours and can reach an apex of 10^9^ − 10^13^ haploid individuals in the bloodstream (Arrow et al. 2004b; Kappe et al. 2010; Graumans et al. 2020). With each 48-hour replication cycle, less than 10% of the asexual parasites terminally differentiate into sexually competent cells called gametocytes (Carter et al. 2013; Collins et al. 2018; Reuling et al. 2018; Tadesse et al. 2019). Gametocyte differentiation is determined at the level of the schizont to either male or female with each emerging and invading merozoite developing and maturing into a single gametocyte, which takes an additional 9-12 days within the human host (Hawking et al. 1968; Silvestrini et al. 2000; Smith et al. 2000; Vaughan et al. 2012). Mature gametocytes do not undergo further replication, and are the only mode of transmission from a human to a mosquito (Chawla et al. 2021); parasites in the asexual erythrocytic stage die after ingestion by the mosquito (Smith and Jacobs-Lorena 2010).

The mosquito phase of the life cycle, which lasts from ∼11-21 days, begins upon the ingestion of infected blood by a female *Anopheles*, with approximately 1-10^3^ *P. falciparum* gametocytes likely to be ingested, representing an orders of magnitude bottleneck from the blood stage (Kappe et al. 2010; Lin et al. 2014; Sato 2021). Once the gametocytes are exposed to the mosquito midgut environment, they differentiate into sexually functional **gametes** (Figure 1; bold text refers to terms shown in the figure). The gametes fuse to form a diploid zygote. The zygote immediately undergoes meiosis and differentiates into an **ookinete** (Figure 1), which then has four haploid genomes in its nucleus (*i.e*., has a ploidy of 4*N*) for ∼17-23 hours (Mzilahowa et al. 2007). The motile ookinete journeys towards the midgut lining, where ∼80% of them are destroyed by the mosquito immune system (Shiao et al. 2006). The motile ookinete forms an **oocyst** (Figure 1) that is embedded on the outer midgut lining of the mosquito and undergoes several rounds of mitosis (Graumans et al. 2020). Once fully matured (taking 11-16 days post-blood meal), the oocyst ruptures and releases 3 × 10^3^ − 2 × 10^4^ haploid sporozoites that invade the salivary gland of the mosquito (Rosenberg and Rungsiwongse 1991; Smith et al. 2014; Wang et al. 2018; Musiime et al. 2019; Kanatani et al. 2024). A severe bottleneck occurs at transmission, where only approximately 20% of sporozoites reach the **salivary glands** where they are stored until less than 1% are transmitted during the infectious bite (Figure 1) when the mosquito takes its next bloodmeal (Rosenberg and Rungsiwongse 1991; Graumans et al. 2020).

The duration of the parasite’s development is limited by the mosquito’s feeding behavior and lifecycle., After mating, female *Anopheles* mosquitoes immediately seek a blood meal and undergo a gonotrophic cycle (*i.e*. pregnancy) for ∼3 days before seeking another host as a potentially infective vector. (Chavasse 2002). Adult female *Anophele*s mosquitoes are generally short-lived, and typically survive for ∼1-3 weeks in optimal conditions (Arrow et al. 2004b; Lambert et al. 2022). While *Anopheles* vectors such as *A. arabensis* and *A. stephensi* tend to take one blood meal per gonotrophic cycle, major malaria vectors such as *Anopheles gambiae* and *Anopheles funestus* frequently take multiple blood meals per gonotrophic cycle (Scott and Takken 2012), with the feeding intervals ranging from 2-4 days. In sum, these constraints imply that *P. falciparum* may spend ∼11-21 days in the mosquito host before transmission to a human host. This varies across vector species and ecological settings, reflecting variation in the parasite’s rate of development, the mosquito’s lifespan, and feeding behavior (Guelbéogo et al. 2018; Brackney et al. 2021).

Population sizes within the mosquito are generally acquired from lab-reared or field-collected *Anopheles* that acquire infections via membrane or skin feeding assays using rodent models (*e.g.,* Vaughan et al. 2012; Kanatani et al. 2024), artificial feeds from field samples (*e.g.,* Andolina et al. 2024), or natural infections in lab-reared or field-collected mosquitoes. Although there are more than 40 common vectors of *P. falciparum*, estimates from laboratory systems are generally limited to the primary vectors such as *An. gambiae* (*e.g.,* Gouagna et al. 1998) and *An. stephensi* (*e.g.,* Churcher et al. 2012; Wang et al. 2018; Andolina et al. 2024; Kanatani et al. 2024). In contrast, field-collected mosquitoes offer a broader range of species (*An. dirus*, (*e.g.,* Rosenberg and Rungsiwongse 1991); *An. coluzzii*, (*e.g.,* Andolina et al. 2024); *An. arabensis* (*e.g.,* Tadesse et al. 2018). Accordingly, parameter estimates may vary across natural infections and laboratory systems, and our consensus estimates are synthesized across experimental systems, field studies, and prior reviews to provide a plausible range. Additional information on the variation across studies can be found in Supplementary Table 1.

Similarly, the variance in parameters characterizing the life cycle can be substantial (summarized in Supplementary Table 1). For one, *P. falciparum* infections are frequently asymptomatic. Most community surveys reveal a higher proportion of asymptomatic than symptomatic infections, even in lower transmission settings (73-98%; as reviewed in Lindblade et al. 2013). Due to immunity in asymptomatic hosts, the parasite population generally reaches a smaller carrying capacity of 10^6^-10^8^ individuals (Tadesse et al. 2018), as opposed to symptomatic infections that span 10^8^- 10^12^ individuals (Tadesse et al. 2018; Uyoga et al. 2021). Similarly, asymptomatic infections last longer: ∼1-2 months, but can persist for more than a year (Briggs et al. 2020; reviewed in Hailemeskel et al. 2024). In contrast, symptomatic infections are generally truncated by treatment to last ∼3-7 days but without treatment can transition to asymptomatic infections (White 2017). Secondly, human hosts remain infectious beyond asexual parasite clearance. Gametocyte clearance was known to persist for months prior to the widespread implementation of ACT. ACT treatment significantly reduces the infectious window to <14 days in most cases (Yilma et al. 2025). Asymptomatic infections may sustain gametocyte carriage for several weeks (Tadesse et al. 2018). Thus, the parasite’s life cycle can be highly variable.

A detailed literature survey of previous estimates of the approximate within-host population sizes, the duration of the different stages of their life cycles, and the corresponding ploidy (Supplementary Table 1, Figure 1), points us to a few key observations. Firstly, *P. falciparum* exists predominantly in a haploid state (∼97% of its life cycle), while it is diploid and tetraploid for only a small fraction (∼3%) of the time (Supplementary Table 2). Thus, heterozygous effects of selected mutations are important for a relatively limited amount of time during their life cycle, and therefore parasite populations can be modeled as haploid. While population-genetic dynamics are very similar in a haploid and diploid population under neutrality, and/or when selected mutations are semi-dominant (*i.e*., the two alleles contribute equally to fitness), this is not necessarily the case with recessive/dominant selected mutations. In particular, when present in diploids or polyploids, partial or fully recessive mutations have reduced or absent phenotypes. Thus, recessive deleterious mutations with full phenotypes in haploid populations can be more effectively selected against, reducing the expected genetic load (*i.e*., reduced mean fitness of the population compared to that of a mutant-free population) that haploid populations carry (reviewed in Otto and Gerstein 2008). In addition, assuming the same number of individuals (*N*) in a haploid and diploid population, if the population-scaled rate of new beneficial mutations (*u*_b_) is low (*i.e*., *Nu*_b_<<1), the rate of fixation of beneficial mutations in haploid populations can be much larger (or smaller) than the diploid population if the beneficial mutation is recessive (or dominant; Orr and Otto 1994). On the other hand, if beneficial mutations are common (*i.e*., *Nu*_b_ ≥ 1), haploid populations will always fix beneficial mutations at higher rates. Therefore, haploidy is central to accurately modeling the dynamics of selected mutations in *Plasmodium* populations and expectations from diploid populations may not naturally apply to *P. falciparum* populations.

Secondly, there are at least 3 bottleneck events during one lifecycle, with the most severe being ∼10^6^-fold at the point of human to mosquito transmission. Moreover, this event follows an extremely rapid exponential growth in the human blood. Recurrent bottlenecks can lead to multiple merger events (Birkner et al. 2009; Tellier and Lemaire 2014), which refers to the coalescence of more than two lineages in the same generation. Moreover, recent experimental evidence has indicated that some strains are more likely to proliferate in the mosquito than others (Li et al. 2019), possibly leading to a wider distribution of offspring numbers than assumed in a Wright-Fisher population (referred to as progeny skew). This suggests that multiple merger events are highly likely during within-host evolution of *P. falciparum*. In other words, when reconstructing the genealogy backwards in time, multiple lineages are likely to coalesce into one at the same node, suggesting that multiple merger coalescents (Pitman 1999; Möhle and Sagitov 2001; Möhle and Sagitov 2001) might best model their coalescent genealogy as opposed to the Kingman coalescent (Kingman 1982), used most commonly.

In general, multiple merger events result in a U-shaped site frequency spectrum (Eldon and Wakeley 2006; Blath et al. 2016), lead to an increase in linkage disequilibrium between alleles (Eldon and Wakeley 2008), and an increase in differentiation between populations (measured by *F*_ST_; Eldon and Wakeley 2009), thus affecting expectations of various population genetics summary statistics. While it is clear that multiple merger events will be common within the mosquito and human hosts, it is unclear whether such events during recurring bottlenecks will affect the genealogy at longer time scales (Tellier and Lemaire 2014; Irwin et al. 2016). If the number of hosts is large and the sampling is geographically scattered and species-wide, the long-term genealogy may converge to the Kingman coalescent (Wakeley and Aliacar 2001; Städler et al. 2009; Heuer and Sturm 2013). However, this is an important unexplored question for the future, which will have consequences for modeling *P. falciparum* populations accurately, and the answer is likely to depend on the transmission dynamics and the sampling scheme (see section *Towards Building a Baseline Model*).

### MUTATION RATE

As *de novo* mutations are the primary source of genetic variation, quantifying the rate at which new mutations arise is an essential component in understanding population genetics in *Plasmodium* populations. While the rate at which new mutations enter *P. falciparum* populations varies across life cycles and time periods (Table 1), the 48-hour asexual erythrocytic cycles are the most important for replication-based mutations. During the asexual erythrocytic cycles, the haploid parasite rapidly replicates in human red blood cells, reaching extremely high within-host population sizes (Figure 2), which vastly increases the possible points of origin for new mutations to arise. Mutation accumulation experiments (Kibota and Lynch 1996; reviewed in Halligan and Keightley 2009), where a single individual is propagated in the laboratory at ideally a population size of 1, thus maximizing the effects of genetic drift, are used to estimate the rate of *de novo* mutations. Such studies in *P. falciparum* (where lab cultures had small population sizes) have indicated that the rate at which new single base substitutions are observed over the 48-hour mitotically dividing period, the asexual erythrocytic cycle, is between 2.10 × 10^−10^ and 5.23 × 10^−10^ per site/ 48 hours (Bopp et al. 2013; Claessens et al. 2014; Hamilton et al. 2016; McDew-White et al. 2019; Table 1). However, strongly deleterious mutations that arise may not be easily observed by mutation accumulation experiments as they are purged from the population prior to sampling (Eyre-Walker and Keightley 2007). The observed mutation rate of single-base substitutions in these studies was adjusted to account for unobserved mutations of strong deleterious effect based on the expected and observed ratios of nonsynonymous and synonymous base substitutions compared to the reference genome (3D7) to approximate the true *de novo* mutation rate This mean adjusted base substitution rate accounting for strongly deleterious mutations ranges from 0.854 × 10^−9^(McDew-White et al. 2019) to 1.70 × 10^−9^ (Bopp et al. 2013) per site per asexual erythrocytic cycle. Assuming 5 mitotic divisions per 48-hour cycle (McDew-White et al. 2019), we may therefore expect the adjusted per-mitotic division mutation rate to range from 1.71 × 10^−10^ (McDew-White et al. 2019) to 3.40 × 10^−10^ (Bopp et al. 2013).

**Figure 2.**
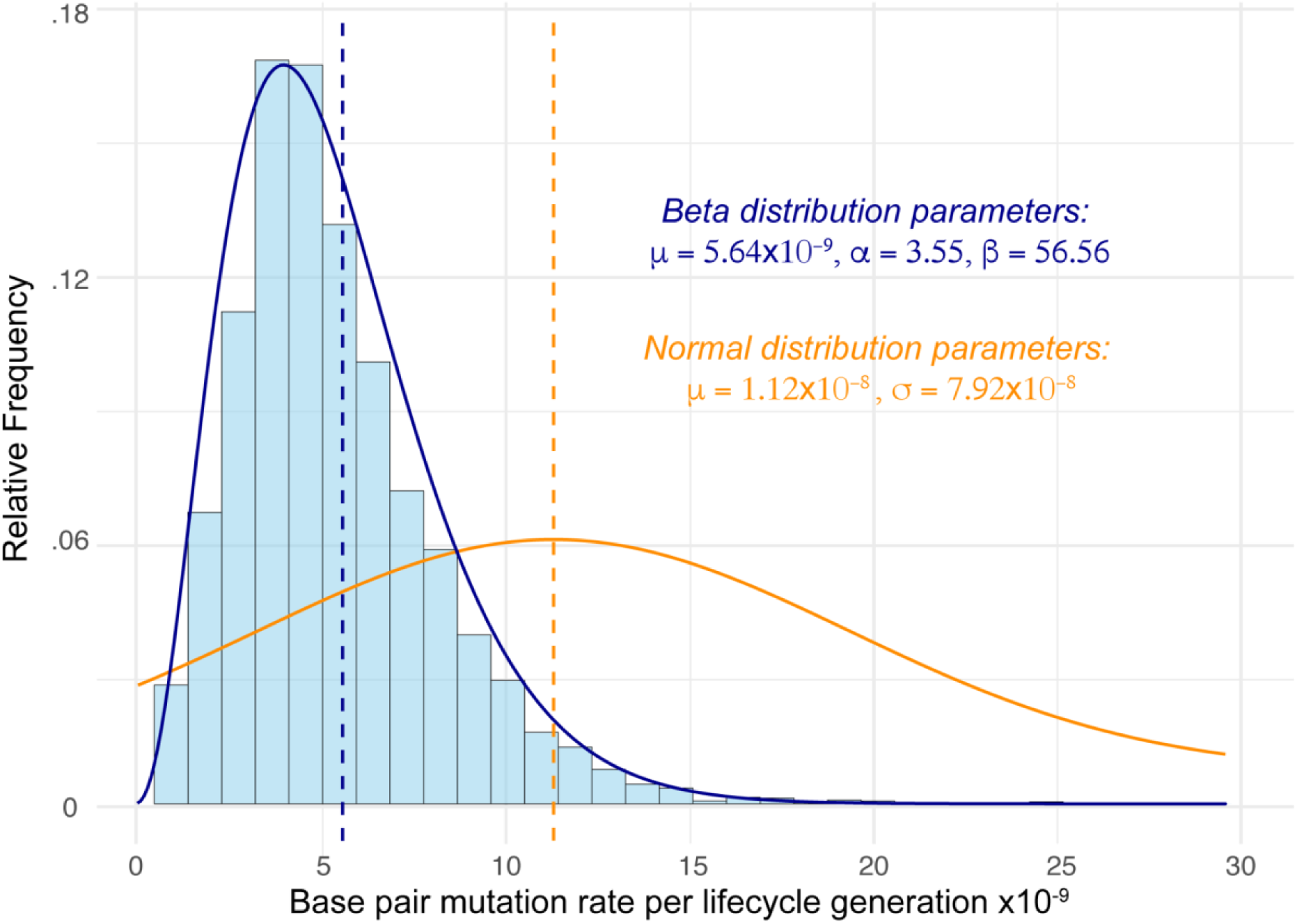
Modeling mutation rate heterogeneity across the *P. falciparum* genome. The histogram displays per-gene synonymous divergence rates (*K_s_*) between *P. falciparum* and *P. reichenowi* directly estimated for 4931 genome-wide genes (available as Supplementary Data 4 in Otto et al. 2014). The dark blue line represents the beta distribution fitted to the histogram data, linearly rescaled around the mean per site per generation single base substitution rate calculated from the mutation rate estimated during the48-hour erythrocytic asexual cycle by McDew-White *et al*. (2019). The orange line shows the commonly used parameters fitted to a normal distribution for the per site per 48-hour erythrocytic asexual cycle mutation rate estimated by Bopp *et al*. (2013). Both estimates are adjusted for strongly deleterious and lethal mutations and assume 33 mitotic divisions per generation. Mean values for each distribution are indicated with dashed lines. We suggest that mutation rate heterogeneity in *P. falciparum* be modeled using the beta distribution (in blue) rather than the normal distribution (in orange).

**Table 1.** Estimated *P. falciparum de novo* mutation rates for single bases (per site) and microsatellite repeats (per locus) over specified periods. Here, “Asexual cycle” refers to a 48-hour mitotically dividing period corresponding to the asexual blood stage in Figure 1. All methods were performed on experimental populations except for longitudinal sampling by Chenet et al. (2015), which involved repeated sampling of a wild Colombian population. Confidence intervals (95%) are provided where available.

| Annotation | Rate | Period | Method | Reference |
| --- | --- | --- | --- | --- |
| Single base substitutions | $3.18 \times 10^{-10}$<br>($2.44 \times 10^{-10} - 3.92 \times 10^{-10}$) | Asexual cycle | Mutation accumulation<br>(31 clones) | McDew-White et al. 2019 |
| | $5.23 \times 10^{-10}$<br>($3.69 \times 10^{-10} - 6.77 \times 10^{-10}$) | Asexual cycle | Mutation accumulation<br>(3 clones) | Bopp et al. 2013 |
| | $2.10 \times 10^{-10}$<br>($1.49 \times 10^{-10} - 2.72 \times 10^{-10}$) | Asexual cycle | Mutation accumulation<br>(37 clones) | (Hamilton et al. 2017) |
| | $4.07 \times 10^{-10}$ (CI not available) | Asexual cycle | Mutation accumulation<br>(19 clones) | Claessens et al. 2014 |
| Microsatellites | $4.45 \times 10^{-7}$<br>($3.48 \times 10^{-7} - 5.42 \times 10^{-7}$) | Asexual cycle | Mutation accumulation<br>(37 clones) | Hamilton et al. 2016 |
| | $4.43 \times 10^{-7}$<br>($4.06 \times 10^{-7} - 4.81 \times 10^{-7}$) | Asexual cycle | Mutation accumulation<br>(31 clones) | McDew-White et al. 2019 |
| | $1.59 \times 10^{-4}$<br>($6.98 \times 10^{-5} - 3.70 \times 10^{-4}$) | Meiosis | Genetic cross<br>(35 progeny) | Su et al. 1999;<br>Anderson et al. 2000 |
| | $0.54 - 3.77 \times 10^{-2}$<br>($8.26 \times 10^{-4} - 7.37 \times 10^{-2}$) | Per month | Longitudinal sampling<br>(79 haplotypes) | Chenet et al. 2015 |

Although all three studies that measured mutation rates in in-vitro experimental populations of *P. falciparum* agree on the final estimated mutation rates, Hamilton et al. (2017) observed fewer indels that are not multiples of three (*i.e.*, indels that result in a frameshift) in coding regions than expected, suggesting that the effect of purifying selection was present in these experimental populations. It is thus highly likely that the estimated rates of *de novo* mutations are presently underestimated, especially because it is not straightforward to account for lethal mutations in mutation accumulation experiments. We can also estimate how many mitotic asexual divisions are likely in a single generation (*i.e.*, between their sexual cycles).

There are ∼3-7 divisions of the zygote in the mosquito gut, ∼13-14 divisions in the human liver, and ∼14 or more divisions in the human blood, totaling ∼30-35 divisions during the life cycle (see Figure 1). Assuming the rate of mutation per cell division estimated by McDew-White et al., and that the parasite undergoes on average 33 cell divisions in a generation, the rate of *de novo* mutations of single base substitutions would be *5*.*6* × *10*^−*9*^ per site/ generation, somewhat higher than previously believed. Mutation rates in unicellular eukaryotes (∼10^-9^-10^-11^ per site/generation) are typically similar to those of eubacteria and much lower than in multicellular eukaryotes (∼10^-8^ – 10^-9^ per site/generation; Lynch et al. 2016). While the *P. falciparum* mutation rate seems to be on the higher end of the range observed in unicellular eukaryotes, with most free-living species like *Chlamydomonas reinhardtii* (3.8×10^-10^ per site/gen), *Paramecium tetraurelia* (1.9×10^-11^ per site/gen) and *S. cerevisiae* (2.6×10^-10^ per site/gen) having much lower rates, the vector of African trypanosomiasis, *Trypanosoma brucei* (1.4×10^-9^ per site/gen) and the fungus *Neurospora crassa* (4.1×10^-9^ per site/gen) exhibit higher mutation rates, very similar in magnitude to *P. falciparum* (Lynch et al. 2016). Moreover, the number of mitotic cell divisions in the human blood can be as high as 40 (see Supplementary note A), giving a maximum of 61 divisions in total during their life cycle, resulting in an even larger upper bound of mutation rate: 1.04×10^-8^ per site/gen.

Mutation rate exhibits heterogeneity across genomic landscapes, influenced by factors such as nucleotide composition, local sequence context, and methylation status (Nachman and Crowell 2000; Ness et al. 2015; Lynch et al. 2016; Smith et al. 2018). While region-specific rates may not have been directly estimated for *P. falciparum*, the distribution of mutation rates across the genome is expected to follow rates of fixed differences at putatively neutral synonymous sites (*K_s_*) between closely related species. Thus, by fitting a beta distribution (*α* = 3.55, *β* = 56.56; *SE* = 0.07, 1.18) to the genome-wide per-gene *K_s_* values between *P. falciparum* and *P. reichenowi* (closely-related species causing malaria in chimpanzees and gorillas) reported by Otto et al. (2014), we obtain an approximate distribution of the mutation rate across the genome which can be scaled around a mean estimate of choice, such as the adjusted rate of 5.6 × 10^−9^ per site/generation estimated above (Figure 2). This distribution will allow researchers to model mutation rate heterogeneity across the *Plasmodium* genome.

*P. falciparum* has an extremely high AT content (80.6% of sites genome-wide) compared to other eukaryotes and *Plasmodium* species (Hamilton et al. 2017) across a highly repetitive genome (Zilversmit et al. 2010). This leads to high genome-wide rates of *de novo* small tandem repeat indels, *i.e.*, microsatellites, due to replication slippage (Lovett 2004), prevalent in both coding and noncoding regions (Hamilton et al. 2017). Microsatellite indel rates observed per 48-hour erythrocytic asexual cycle from Hamilton et al. (2017) and McDew-White et al. (2019) were highly consistent (Table 1). In addition, studies have estimated microsatellite indel rates observed between meiotic generations (Su et al. 1999; Anderson et al. 2000), as well as per month across a wild Colombian population using a Bayesian phylogenetic approach (Chenet et al. 2015), which could be useful if *de novo* mutations were likely to accrue as a function of time rather than the number of cell divisions (Gao et al. 2016). However, microsatellite indels that are not multiples of three are disproportionately uncommon in coding regions compared to intron/intergenic regions due to strong purifying selection against frameshift *de novo* microsatellite mutations. As a result, the values reported in Table 1 represent the lower bounds of the *de novo* microsatellite mutation rates, while the true rates may be up to 48% higher when accounting for unobserved lethal or strongly deleterious frameshift mutations (Hamilton et al. 2017).

### GENOME ARCHITECTURE

The AT-rich 23 Mb *P. falciparum* genome is dense with coding regions, with exonic sites of over 5267 genes covering approximately 52% of the genome; the mean length of exons, introns and intergenic distances is 949 bp, 179 bp, and 1,694 bp respectively, with an average (and median) of 2.39 (and 1) exons per gene with 47% of genes identified as having no introns (details available in Gardner et al. 2002). In eukaryotes with similar genome-wide coding sequence density, such as *Drosophila melanogaster* (dos Santos et al. 2015), this would be expected to result in pervasive linked effects of selection (Begun and Aquadro 1992; Charlesworth et al. 1993; Comeron et al. 2012). These estimates notably exclude any extrachromosomal DNA and the highly variable subtelomeric regions that contain major gene families responsible for immune evasion and virulence, such as *Var* genes (Otto, Böhme, et al. 2018). Because these regions are difficult to align and find orthologous regions in (Otto, Böhme, et al. 2018), they are routinely excluded from population genetics studies.

Comparative genomics analysis has uncovered that approximately 60% of *P. falciparum* genes are conserved among all five *Plasmodium* species studied (Cai et al. 2012); this set of genes likely represents conserved essential functions across the entire genus. However, note that due to ascertainment bias, predicted gene sequences from many protists, especially those that have diverged substantially from Opisthokonts (*e.g.*, those belonging to Alveolates, Discobids), lack known orthologs and assigned functions (Burger et al. 2016; Johri et al. 2019; Prieto-Baños et al. 2025). Thus, as expected, when sequence-independent computational algorithms based on geometry were applied to the unannotated set of genes in *P. falciparum*, similarity to previously known protein domains was identified in 25% of these genes (Behrens and Spielmann 2024), suggesting that these represent putatively functional coding regions.

Due to the high AT-content of the *P. falciparum* genome, fewer than a third of coding variant sites are synonymous; we estimate the proportion to be 17.5%. Thus, assuming that mutations at all nonsynonymous sites have some effect on the fitness of the individual, the proportion of coding sites experiencing direct selection would be 82.5%. The proportion of noncoding regions under purifying selection is unclear. However, previous studies (Siepel et al. 2005) that have used phylogenetic methods to estimate this proportion across model organisms found that in the human, *C. elegans*, *D. melanogaster*, and *S. cerevisiae* genomes where ∼3%, 30%, 18%, and 70% are annotated as coding, ∼ 2%, 11%, 20%, and 6% of their genomes were predicted to be conserved noncoding respectively. As the *P. falciparum* genome resembles the *S. cerevisiae* genome in terms of the coding density (∼52.6%), total number of genes (∼5k vs ∼6k in yeast) and the genome size (23 Mb vs 12 Mb in yeast), we assume that ∼5-10% of their noncoding regions are conserved. As such, assuming that 82.5% of coding regions in *Plasmodium* genomes are conserved (43% of genome-wide sites) and ∼10% of genomic sites represent conserved noncoding regions, approximately 53% of genome-wide sites likely experience direct selection. Further analysis is needed to more accurately identify and quantify phylogenetically conserved sites with approaches such as phastCons (Siepel et al. 2005) or GERP (Davydov et al. 2010), which in turn would enable analyses to effectively quantify the direct and linked effects of selection genome-wide. There is a striking conservation of the number of chromosomes, number of genes, and synteny across the seven species belonging to the Laverania genus, which includes *P. praefalciparum* and *P. reichnowi*, and reasonable genomic synteny extends to *P. ovale spp.* (Rutledge et al. 2017), as well as to *P. vivax*, *P. knowlesi*, and *P.y. yoelli* that share ∼77% of orthologs with *P. falciparum* (Carlton et al. 2008). With whole genome sequences of 14-15 closely related taxa, such an endeavor seems plausible.

### RECOMBINATION AND SELFING RATE

Recombination refers to the shuffling of genetic variation that occurs during meiosis, where genetic material from two homologous chromosomes is exchanged. Recombination results in novel combinations of genetic variants, ultimately bolstering the effectiveness of natural selection. Due to its influential role in shaping patterns of genomic variation, understanding the heterogeneity in recombination rate across the genome and between populations is important to understand the patterns of genetic diversity in a species (Cutter and Payseur 2013; Peñalba and Wolf 2020). *P. falciparum* undergoes sexual recombination, between the chromosomes from a haploid female macrogamete and a haploid male microgamete, once per life cycle in the mosquito (Figure 1) (Mzilahowa et al. 2007). Numerous studies have characterized the patterns of recombination in *P. falciparum* using microsatellite markers or tiling arrays (Walliker et al. 1987; Walker-Jonah et al. 1992; Kerr et al. 1994; Su et al. 1999; Hayton et al. 2008; Jiang et al. 2011). Recent work by Miles et al. (2016) used short-read whole genome sequencing to achieve a comprehensive understanding of recombination in *P. falciparum* crosses by sequencing the parents and offspring from three pairs of crosses between diverse lab strains. This study achieved a resolution of ∼300 bp for each recombination event using SNPs and indels in the strains, which was an order of magnitude greater than previous work using an SNP array (Jiang et al. 2011). The average recombination rate for all three crosses was 7.4 × 10^-7^ crossovers/bp per meiosis, confirming the previous observation of a recombination rate that is an order of magnitude higher than mammals such as rats, mice, and humans (Jensen-Seaman et al. 2004) but smaller than an estimate of 3.5 × 10^-6^ in yeast (Ruderfer et al. 2006). More recent genetic crosses generated in human liver-chimeric mice produced similarly high map lengths and recombination rates, indicating that elevated recombination is a consistent feature across *P. falciparum* crosses (Button-Simons et al. 2021). Because unicellular eukaryotes generally have small chromosomes and at least one crossover is required per meiosis, this results in a relatively high rate of crossover on average (Lynch 2006).

The probability of gene conversion per base pair (per haploid genome) was estimated to be 3.6 × 10^-7^ per meiosis, with average tract lengths of 1.4 kb, after adjusting for small gene conversion events that were likely missed. These gene conversion tract lengths are also similar to those of yeast (Mancera et al. 2008), but longer than those of humans (Jeffreys and May 2004). Genome-wide patterns of recombination showed a significantly lower recombination rate near centromeres and subtelomeres, and with higher recombination rates associated with repetitive DNA. Both recombination and gene conversion showed elevated rates in exons, unlike humans, where recombination primarily occurs in hotspots near but not within the coding regions of genes (Myers et al. 2005). The higher GC content of coding regions in *P. falciparum* may partially explain this preference, but it may also be explained by a methodological bias against finding recombination events in AT-rich noncoding regions. Additionally, a 12-bp motif was found to be significantly enriched for recombination in multiple studies (Jiang et al. 2011; Miles et al. 2016); however, limited data currently prevents the confident classification of any recombination hotspots (Miles et al. 2016). The *prdm9* gene, a well-known mediator of recombination hotspots in many species (Baudat et al. 2010) has a putative ortholog in the *P. falciparum* genome (Jiang et al. 2011), so future studies using a higher number of crosses or a population genetic method to estimate the recombination rate (reviewed in Peñalba and Wolf 2020) may be able to more confidently identify recombination hotspots in the *Plasmodium* genome.

While the actual rate of recombination is quite high, the effective rate of recombination is substantially lower in *P. falciparum* populations. This is mostly because of two reasons: (1) as the female and male gametocytes mature from the same pool of parasite, selfing can occur between genetically identical clones of *P. falciparum*, making recombination ineffective. (2) Due to the bottlenecks during ingestion by mosquitoes in the *P. falciparum* lifecycle, only a small number of strains (usually between 1-20; Nkhoma et al. 2020) are available to recombine. Moreover, the differentiation and pairing of gametes occurs within hours of ingestion by a mosquito (Baton and Ranford-Cartwright 2005), so recombination events usually occur between parasites ingested from a single human host rather than strains from multiple blood meals (Camponovo et al. 2023). This is likely to increase the probability of selfing and mating between genetically highly related chromosomes, reducing the effective rate of recombination in the population.

While the rate of selfing is likely to be non-zero, it will depend on the number of distinct strains in the mosquito. For instance, assuming random mating, the rate of selfing with 10 distinct haplotypes will be 0.1, while it will be 0.5 with 2 distinct haplotypes. Thus, the effective rate of recombination will mostly depend on the overall level of inbreeding occurring in the population, *i.e*., *R_eff_* = *R*(*1* − *F_it_*). Here *R* is the population-scaled rate of recombination, and *F_it_* represents the overall level of inbreeding, which will be influenced by both selfing events and mating between closely related individuals, and is calculated by comparing the heterozygosity of parasite populations within the guts of individual mosquitoes (*H_W_*) to that of the entire population of parasites sampled from mosquitoes (*H_S_*) with 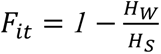 (Manske et al. 2012). As expected, the number of genetically distinct malaria strains present within a human host, called the complexity of infection (COI; Viriyakosol et al. 1995; Snounou and Beck 1998; Paschalidis et al. 2023), was found to positively correlate with the amount of transmission and therefore with rates of outcrossing (Camponovo et al. 2023), leading to higher nucleotide diversity and less linkage disequilibrium in such populations (Anderson et al. 2000).

Several attempts have been made to estimate the inbreeding rate in *P. falciparum* populations using either the parasite population in the mosquito or the human host. Obtaining estimates of inbreeding from infected mosquitoes is the most direct way. Analysis from the mosquito stage is also most relevant to its effect on recombination, particularly because haploid oocysts attached to the mosquito midgut contain thousands of cells descended from one zygote (a single meiosis), and thus can be used to infer the genotypes of zygotes and infer *F_it_* (Ranford-Cartwright et al. 1991). Genotyping of microsatellite loci in oocytes from mosquito midguts in high-transmission areas has yielded a few estimates of inbreeding rate: 0.56-0.6 in Malawi (Mzilahowa et al. 2007), 0.33 in Tanzania (Hill et al. 1995), and 0.4 in Western Kenya (Razakandrainibe et al. 2005). Due to the recent large-scale malaria sequencing efforts from at least 20,000 patient blood samples (Abdel Hamid et al. 2023), inbreeding estimates using *Plasmodium* samples from infected human patients (commonly referred to as *F_WS_*) have become prevalent and are largely correlated with estimates using COI, though they use a relatively indirect approach (Auburn et al. 2012; Manske et al. 2012).

Inbreeding rates obtained using parasite populations in the human host yield somewhat higher estimates from ∼0.9 in African populations to >0.95 in South America, Southeast Asia, and Oceania (Ahouidi et al. 2021; Abdel Hamid et al. 2023; Abdel Hamid et al. 2025) as summarized in Supplementary Table 3. However, a more fine-grained analysis of African *P. falciparum* populations found large amounts of variation in mean *F_WS_*, with countries with high rates of transmission like Kenya and Malawi (*F_WS_* ∼*0*.*7*) having lower rates of inbreeding than countries with low rates of transmission like Senegal and Ethiopia (*F_WS_* > *0*.*95*; Amambua-Ngwa et al. 2019). Thus, it is possible that differences between the mosquito-based and human-based studies simply reflect the differences in the specific populations used - *i.e.*, rates of inbreeding could vary locally vs globally. However, other methodological differences might also contribute to this discrepancy. For instance, estimates using parasites sampled from humans do not account for the drastic bottleneck experienced by the parasites immediately before recombination, and while mosquito-based studies used a few microsatellite loci, the human-based studies have included data from the whole genome. In either case, inbreeding rates seem to vary between 0.4-0.95, governed by rates of transmission in the specific local population.

Because the effective rate of recombination in *P. falciparum* populations depends on the rate of inbreeding, which in turn depends on the rate of transmission, it varies across populations according to the transmission dynamics. The bite frequency and infection rate vary by geographic region. Across multiple studies, there is a consistent pattern of intense hotspots in rural and peri-urban regions, and lower intensity within city centers (Duchemin et al. 2003; Doumbe-Belisse et al. 2018). This is because the number of infected bites per human per year (EIR) is much lower in indoor (31.14) than outdoor (66.65) settings (Doumbe-Belisse et al. 2018). Similarly, across multiple regions in sub-Saharan Africa, the EIR was found to be 7.1 in the city centers, 45.8 in peri-urban areas, and 167.7 in rural areas (Duchemin et al. 2003). Thus, the likelihood of transmission can vary substantially depending on spatial factors and population density.

High-transmission regions in Africa and South Asia have higher nucleotide diversity, and much faster decay of linkage disequilibrium (LD) over distance relative to low-transmission regions in South America, Southeast Asia, and Oceania (Achidi et al. 2008; Ahouidi et al. 2021; Abdel Hamid et al. 2023; Abdel Hamid et al. 2025). However, the magnitude of the difference in LD decay is moderate across populations, with nucleotide diversity in coding regions varying from a median of 0.00019 in Eastern Southeast Asia to 0.00028 in West Africa, and *r*^2^ in coding regions decaying to baseline levels within ∼2,000-10,000 bp for Asian populations and ∼500-1000 bp for African populations (Baudat et al. 2010; Ahouidi et al. 2021). Thus, LD decays in *P. falciparum* much faster than in human populations (Auton et al. 2015) but is similar to *Drosophila* populations, where LD decays in ∼500-10000 bases.

While these estimates are derived from samples pooled from multiple populations collected at different time points, they provide time and space-averaged estimates that may not necessarily correspond to present estimates in local populations. For instance, in regions of varying transmission intensity in Colombia, where clonal genotypes have been observed to persist for up to 16 years(Carrasquilla et al. 2022), LD decay was found to vary drastically between subpopulations, with LD in subpopulations with low transmission decaying to background levels in ∼500 kb and relatively higher transmission areas within ∼250 kb (Echeverry et al. 2013). Similarly, certain malaria outbreaks in low transmission regions like eastern Panama (Obaldia et al. 2015) and northwest Ecuador (Sáenz et al. 2015) have been composed of only one to three strains, and high levels of monoclonal infections have been identified in regions with low transmission like Cambodia (Parobek et al. 2016) or the highlands of Ethiopia (Holzschuh et al. 2024). The LD in Cambodian populations of *Plasmodium* has been shown to decay to baseline levels hundreds of base pairs slower than populations in sub-Saharan Africa, supporting reduced recombination in low transmission regions (Miotto et al. 2013; Samad et al. 2015). Studies using identity-by-descent (IBD) tracts support similar observations (*e.g.*, Taylor et al. 2017; Osborne et al. 2024), while offering finer spatial resolution because they directly capture recent shared ancestry. Thus, the levels of inbreeding and the effectiveness of recombination varies at the local scale, with regions with high transmission and endemicity generally having more effective recombination than regions with lower or declining malaria transmission (Anderson et al. 2000).

Assuming a Wright-Fisher panmictic population, a previous study estimated the haploid effective population size of *P. falciparum* in Senegal to be 50,000 (Chang et al. 2012). Assuming the relevant value of the per-site rate of recombination and inbreeding estimates of 0.6 and 0.9 for high vs low transmission populations, we find the theoretical expectation of the LD-decay (Figure 3) to match empirical estimates quite well (see Figure 2c of Ahouidi et al. 2021). We theoretically predict LD to reach baseline levels in ∼250 bp in high transmission vs ∼1000 bases in low transmission populations. An important implication of vastly different recombination rates across populations is that the expected signatures of recent selective sweeps would likely be highly population-specific, deeming it necessary to generate population-specific expectations when performing selection scans using the *P. falciparum* genome (see the section *Drug resistance and selective sweeps* for a more detailed discussion).

**Figure 3:**
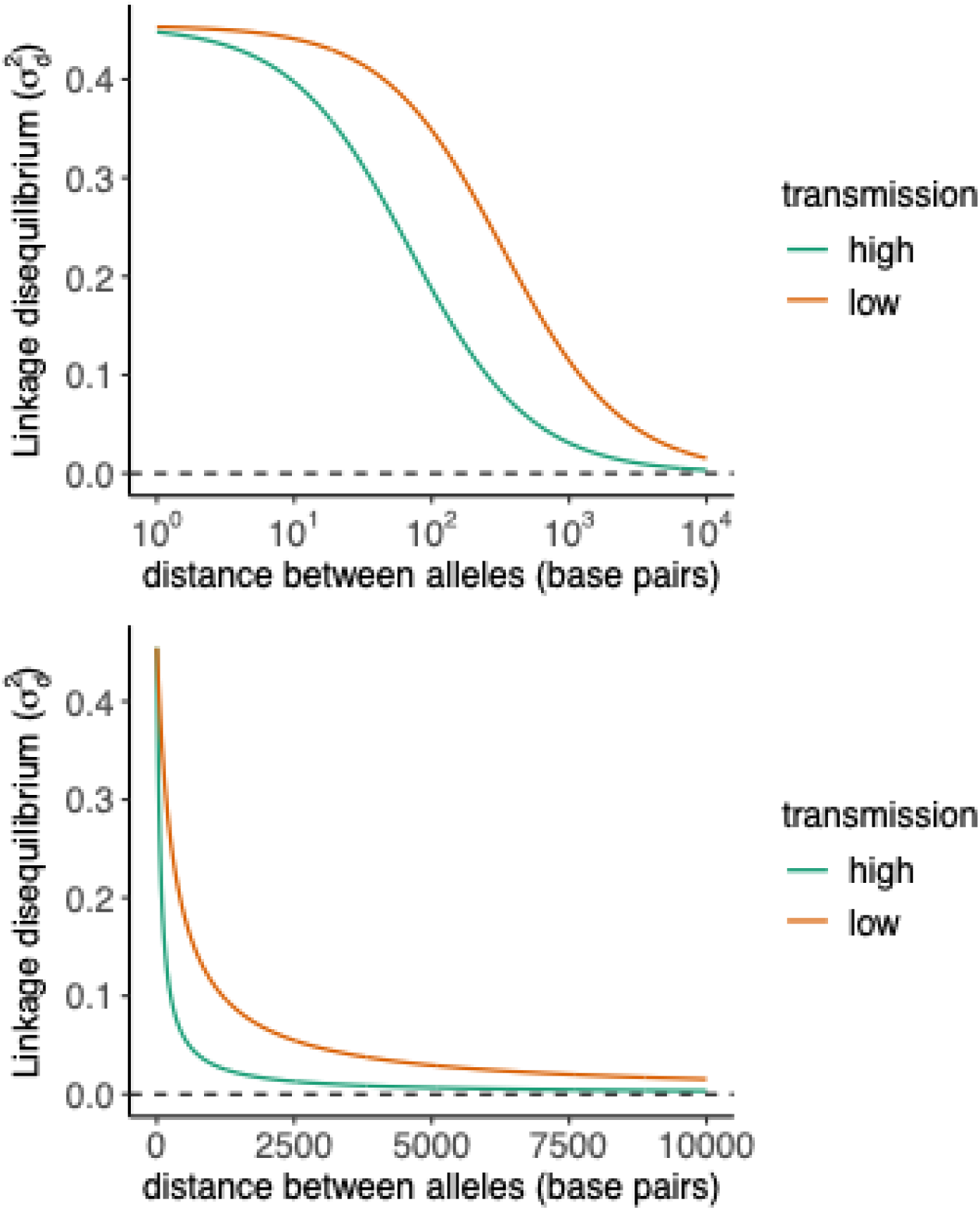
The expected decay of linkage disequilibrium (between hosts) with distance between alleles in a high vs low transmission population of *P. falciparum*. Here, *N*=80,000, *r* = 7.4 × 10^-7^ per site/generation, and *N*_e_=*N*/(1+*F*) where *F*=0.6 and 0.9 in high and low transmission populations respectively. Here, linkage disequilibrium is measured using 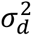such that 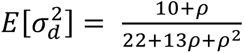, where *ρ* = *2N_e_ rx*(*1* − *F*), and *x* is the distance between alleles (Ohta and Kimura 1971).

Although note that empirical estimates of LD decay in *P. falciparum* have been generated using unphased genomes, by employing the method of Roger and Huff (Rogers and Huff 2009) after filtering and simplifying polyclonal infections. In this method, for each pair of SNPs, a correlation of the unphased diploid genotypes (2/1/0) is used to approximate *r*^2^. In the pooled sequencing of parasites within human hosts, a heterozygote genotype does not represent the presence of the two alleles at 50% frequency, but instead reflects a mixture of haploid parasite strains within the same blood sample. Mixed infections may obscure the true haplotype composition, where unphased genotypes may either mask the association between alleles at different loci, decreasing LD, or cause distinct haplotypes to appear identical, inflating LD. Thus, in high transmission populations where within-host diversity may be substantial, estimates of LD decay may be biased. In samples from monoclonal infections, the LD decay curve is more likely to be accurate.

### DISTRIBUTION OF FITNESS EFFECTS

New *de novo* mutations occur every generation (see the section *Mutation Rate*) and can either be harmful or beneficial for the individual, have the same fitness effect as the wildtype, or may have no effect on fitness at all. The latter category of mutations is called “neutral” mutations, examples of which are many synonymous mutations and those that occur in intergenic regions that serve no function. Unsurprisingly, representing random changes to a complex system, most new mutations in functionally important regions tend to be deleterious, while a small minority (less than ∼1%) may be beneficial (Eyre-Walker and Keightley 2007; Bank et al. 2014). Although we understand this broad-level classification of mutations, the magnitude of the effect of each mutation on fitness follows a continuous distribution between −1 (signifying lethal) and +1 (signifying the fittest individual). This distribution of fitness effects (DFE) of new mutations is important to understand the expected amount of genetic variation in a population, the amount of genetic load, the rate of Muller’s ratchet, and the effects of selection on linked sites.

Most genes in *P. falciparum* exhibit low values of divergence at nonsynonymous vs synonymous sites, measured by *d*_N_/*d*_S_, with a median of 0.17 (Jeffares et al. 2007) and a mean *d*_S_ value of 0.057 when using *P. reichnowii* (Prugnolle et al. 2008), consistent with widespread purifying selection. Fast-evolving genes tend to have low expression, suggesting that these are likely under reduced selective constraints, consistent with universal trends. However, some of the *P. falciparum* coding regions have been found to comprise lowly conserved domains that exhibit higher levels of nonsynonymous substitutions (*d*_N_/*d*_S_ ∼0.35) and have been predicted to represent low complexity hydrophilic non-globular domains (Wootton 1994; Gardner et al. 1998; Gardner et al. 2011). Gardner et al. (2011) found that for a subset of 40 genes that have conserved orthologs and synteny in the *P. falciparum* genome, these protein domains of low conservation occupy ∼50% of the total coding sequence, suggesting that a substantial number of new mutations in coding regions may be mildly deleterious and/or effectively neutral. Relatedly, the observed amount of polymorphism at nonsynonymous (*π_N_*) vs synonymous sites (*π_S_*), measured by *π_N_*⁄*π_S_* in *P. falciparum* populations, was found to be unexpectedly high, with a mean value of ∼0.52 and with many genes exhibiting values above 1 (Chang et al. 2012). Similarly, the site frequency spectra of both nonsynonymous and synonymous sites were found to be very similar in *P. falciparum* populations (Chang et al. 2012), suggesting the presence of many mildly deleterious mutations.

However, because the mutation spectrum of *de novo* mutations is AT-biased (Hamilton et al. 2017), it is unclear what effect that might have on the expected statistics. Alternatively, it is possible that the unique life cycle of the parasite with multiple bottlenecks results in summary statistics at selected and neutral sites being similar (Chang et al. 2013). Thirdly, the global population of *P. falciparum* may simply not be in equilibrium either due to recent demographic fluctuations or the relatively ancient speciation bottleneck that occurred 40,000-60,000 years (or ∼400,000 generations) ago (Otto, Gilabert, et al. 2018). Because variation at effectively neutral sites takes longer to reach equilibrium levels than at selected sites (Brandvain and Wright 2016), nonequilibrium demography could explain higher diversity at nonsynonymous relative to synonymous sites. Finally, selection on synonymous sites might also yield such observations (Musto et al. 1999; Peixoto et al. 2004; Chan et al. 2017; Sinha and Woodrow 2018). These hypotheses need to be tested methodically to better understand patterns of variation observed in *P. falciparum* populations.

Population genetics methods to estimate the DFE using the SFS have been applied to *P. falciparum* populations (Morgan et al. 2020; Perkins et al. 2025) and have inferred a DFE highly skewed towards mildly deleterious mutations. However, these methods assume a diploid Wright-Fisher population. While these methods have been found to be robust to many model violations in diploid populations, their accuracy has not been tested for haploid organisms or those with complex life cycles and transmission. Moreover, in species that exhibit strong progeny skew, the methods provide highly inaccurate results (see Figure 1 of Morales-Arce et al. 2022). Thus, methods that account for the *P. falciparum* life cycle and transmission are needed to accurately infer the DFE.

While there are no experimental estimates of fitness effects of new mutations in *P. falciparum* yet, it seems like a feasible goal, as both gene editing and in-vitro competition assays have been successfully performed to study drug resistance (*e.g.* Walliker et al. 2005; Stokes et al. 2021; Hagenah et al. 2024). Although such studies need to be performed for more genes, the few mutations evaluated in genes involved in drug resistance show that most mutations are either neutral or deleterious (Stokes et al. 2021), in line with genome-wide expectations. A gene knock-out study (Zhang et al. 2018) conducted in-vitro suggested that approximately 50% of all genes are essential during the asexual erythrocytic stage, suggesting that a substantial fraction of large deletions and frameshift mutations might be lethal or strongly deleterious in *P. falciparum* populations, though this number is probably much larger because this study captured only the blood stage, not the entire lifecycle.

Due to the high AT content in the *P. falciparum* genome, the rate of *de novo* insertions/deletions is relatively high (discussed in section *Mutation Rate*). These insertions and deletions in protein-coding regions that are not multiples of three cause frameshift mutations, typically resulting in truncated or misfolded proteins with strongly deleterious effects due to loss of function. This is consistent with an overrepresentation of fixed indels in coding regions whose lengths are multiples of 3 (Jeffares et al. 2007). Insertions/deletions in noncoding regulatory regions, however, might have only mild selective effects. Thus, the DFE that combines the effects of base substitutions and insertions/deletions in functionally important parts of the genome will likely be bimodal, with a peak at strongly deleterious and another at mildly deleterious mutations. Because of the abundance of insertions/deletions in *P. falciparum*, they likely offer variation that is important for immune evasion or other biological functions, and thus their contribution to evolutionary dynamics might be significant.

As the probability of fixation of very mildly deleterious mutations is appreciable, such mutations can contribute substantially to differences between species and subpopulations. They contribute to outliers when identifying targets of local adaptation via population differentiation tests (Johri, Charlesworth, et al. 2021). Therefore, a null model that incorporates the effect of mildly deleterious mutations will be important for correctly interpreting outliers of selection scans across the *P. falciparum* genome.

### EFFECTS OF SELECTION AT LINKED SITES

#### Background selection

Deleterious mutations are purged from populations via purifying selection, and this leads to a decrease in diversity at functionally important sites in the genome. However, alleles linked to deleterious alleles can also be removed by purifying selection acting on selected sites. This process is called background selection (Charlesworth et al. 1993). While strongly deleterious mutations get purged rapidly from populations, mildly and moderately deleterious mutations can segregate in populations for a longer period of time before being purged, resulting in a distortion of the gene genealogy itself (Charlesworth 2013). Thus, background selection not only reduces neutral diversity, but can also skew the expected site frequency spectrum at neutral sites linked to selected sites (Charlesworth 2013; Nicolaisen and Desai 2013). This distortion of expected summary statistics can lead to the misinference of demographic history when using methods that assume strict neutrality (Ewing and Jensen 2016; Johri, Riall, et al. 2021). In particular, background selection results in a skew of the site frequency spectrum towards low-frequency alleles, leading to a false inference of recent population growth. Despite the pervasive effects of background selection demonstrated across multiple species (Cutter and Payseur 2013), currently only a minority of methods account for the effects of selection on linked sites while estimating parameters of demography (Sheehan and Song 2016; Johri et al. 2020; Johri et al. 2023). Moreover, a recent simulation study suggests that selection in *P. falciparum* might bias inferences of demography and population structure (Guo et al. 2024). It is therefore important to understand the extent of background selection that might be expected in *P. falciparum* populations.

The *P. falciparum* genome is ∼23 Mb, with 14 chromosomes, comprising 52.6% coding, 5.7% intronic, and 53.2% intergenic regions. We estimate that 53% of the *P. falciparum* genome is likely to be under direct selection (see the section on *Genomic Architecture* above). As most new mutations at selected sites are deleterious (Eyre-Walker and Keightley 2007), the number of new deleterious single-base pair mutations in a haploid individual per genome per generation (*U*_base_) can be estimated to be 0.53 × μ × 23Mb, where μ is 5.6 × 10^−9^ per site/generation, which is 0.068. Because there are a significant number of indels in *P. falciparum* that are likely to be harmful, we include them in estimating the genome-wide deleterious mutation rate. McDew-White et al. (2019) estimate an average rate of ∼4.4 × 10^-7^ indels per locus/generation (= 8.8×10^-8^ per locus per cell division). Assuming that there are 123,722 loci in the core genome (as found in McDew-White et al. 2019), we estimate that each individual is likely to have ∼0.05 new insertions/deletions every generation. Note that some indels will have no effects on the fitness of the individual, especially if they occur in non-coding regions. Assuming that 53% of them are selected against, we get *U*_indel_∼0.03. This gives us a total *U* ∼ 0.096, *i.e.*, there are likely 0.10 deleterious mutations per individual/ generation.

Using estimates of cross-over rates for chromosomes (see Figure 3 from Miles et al. 2016), we can estimate the expected nucleotide diversity at neutral sites in the presence of background selection (BGS) relative to strict neutrality, denoted by the symbol *B*, (Nordborg et al. 1996). Note that a value of *B* close to one indicates minimal effects of BGS while a value closer to zero suggests a drastic reduction in diversity due to BGS. We find that *B* for an average-sized chromosome in *P. falciparum* is only ∼0.99 and 0.96 in high (assuming *F*=0.6) and low transmission (assuming *F*=0.9) populations, respectively (discussed in the section *Recombination*). Similar values (*B*∼0.96-0.99) can be estimated for chromosomes of different lengths: chromosome 1, 10, and 14 corresponding to 640 kb (shortest), 1.69 Mb (average), and 3.3 Mb (longest), with the chromosome-wide map length in Morgans (*R*) being ∼0.5, 1, and 2.25, respectively (Supplementary Table 4) and accounting for selfing. In summary, the reduction in diversity due to linked effects alone in *P. falciparum* genomes is unlikely to be much, far smaller than that expected in *D. melanogaster* populations (where *B*∼0.5).

Because *P. falciparum* has many chromosomes (14), strongly deleterious mutations on other chromosomes can also lower diversity at a focal chromosome (Santiago and Caballero 1998; Charlesworth 2012), which is referred to as unlinked BGS. We estimate that these unlinked effects of BGS acting on a chromosome of size 1.6 Mb would result in ∼0.96× the nucleotide diversity relative to under neutrality. Assuming multiplicativity of fitness effects, the combined effect of both linked (caused by deleterious mutations on the same chromosome) and unlinked BGS (caused by deleterious mutations on other chromosomes) will result in *B* ∼ 0.93, which we estimate to be similar across all chromosomes (Supplementary Table 4). Thus, the effects of background selection in lowering nucleotide diversity are expected to be minimal in *P. falciparum,* which is consistent with very short chromosomes and thus high rates of recombination in this unicellular organism. However, note that the effect of alternating sexual and asexual cycles on the expected amount of BGS is not clear. During asexual divisions, because the entire genome is linked, the effects of BGS will be much stronger. Additionally, background selection can skew the site frequency spectrum of neutral alleles (Charlesworth 2013; Nicolaisen and Desai 2013), and the extent of this skew in *P. falciparum* remains to be tested, as it can bias the inference of population history. Note that our calculations assume panmixia and equilibrium, both of which are unlikely to realistically represent *Plasmodium* populations. Thus, further investigation accounting for the parasite life cycle and host-pathogen dynamics is needed to fully understand the effects of background selection on genomic variation in *P. falciparum*.

#### Drug resistance and selective sweeps

Since malaria eradication became a global priority in the 20th century, recurrent emergence of antimalarial-resistant mutations has lead to the failure of first-line treatments. Historically, the majority of the resistance mutations, including those resistant to chloroquine, sulfadoxine-pyrimethamine, mefloquine, and some artemisinin resistant lineages, have first emerged in the low transmission Greater Mekong subregion before spreading to other endemic regions. This repeated pattern suggests that transmission intensity may influence the emergence, spread, and successful establishment of drug resistance. Parasites resistant to chloroquine, the first antimalarial widely implemented in the 1950s, have emerged independently at least four times due to mutations in the *P. falciparum* gene encoding the chloroquine resistance transporter (*pfcrt*; Payne 1987; Wellems and Plowe 2001). However, the major resistant mutation responsible for the widespread proliferation of chloroquine resistance across Africa originated in Southeast Asia. (Payne 1987; Wootton et al. 2002). Despite chloroquine being removed as a first-line drug due to its failure, the presence of chloroquine-resistant mutations remains almost entirely fixed in low-transmission populations like Southeast Asia and South America (Bacon et al. 2009; Griffing et al. 2010; Muhamad et al. 2011; Phompradit et al. 2014). In contrast, the chloroquine-resistant *pfcrt* genotype became almost undetectable in the high transmission Malawi population within twelve years after chloroquine’s removal as a first-line drug (Kublin et al. 2003; Frosch et al. 2014; Takala-Harrison and Laufer 2015).

Resistance to the subsequent first-line antimalarial drug, sulfadoxine-pyrimethamine arises from sequential mutations accumulated in dihydrofolate reductase (*pfdhfr*) and dihydropteroate synthase (*pfdhps*; Cowman et al. 1988; Triglia et al. 1997) genes. These highly resistant haplotypes followed a similar pattern to the widespread accumulation of chloroquine resistance, where the lineage emerged in Southeast Asia, spread to Africa, and emerged independently in South America and Oceania (Mita et al. 2007; Mita et al. 2011). Similar to patterns in chloroquine resistance, highly resistant sulfadoxine-pyrimethamine haplotypes remain prevalent in Southeast Asia and South America despite the removal of sulfadoxine-pyrimethamine as a drug, while resistance remains at heterogenous frequencies across regions in Africa (Mita et al. 2011).

Currently, there is a major concern regarding resistant lineages of the first-line treatment of the 21st century: artemisinin combination therapy (ACT). The first evidence of artemisinin resistance (mediated by nonsynonymous mutations in the *pfkelch13* gene) emerged in western Cambodia in 2008 as treatment failed to quickly or completely clear parasite infections (Noedl et al. 2008). By 2013, resistance and associated K13 propeller mutations has been observed in multiple other Southeast Asian countries, such as Vietnam, Thailand, and Laos, associated with resistance to the long-acting drug piperaquine, with substantial increases in the frequency of resistance alleles from 4% to 63% observed in Cambodia from 2007 to 2013, respectively (Saunders et al. 2014; Amaratunga et al. 2016; Imwong et al. 2017; Thanh et al. 2017; Amato et al. 2018). Despite ACT being an effective treatment, there are numerous signs that resistance is rapidly emerging, facilitated through clonal spread of *pfkelch13* mutations (Parobek et al. 2017) that may be maintained by low transmission. Multiple independent *pfkelch13* and associated partner drug mutations have emerged in African populations as well (*e.g.*, Rosenthal et al. 2024; Brhane et al. 2025; Niaré et al. 2025). *Pfkelch13* mutations appear to have initially expanded in Uganda at a similar speed to emergence in Southeast Asia (Rosenthal et al. 2024). Unlike the case of chloroquine and sulfadoxine-pyrimethamine, differences in the flanking haplotypes suggest that these resistance mutations appear to originate in Africa, instead of having spread from Southeast Asia (Conrad et al. 2023; Niaré et al. 2025). Although there are multiple factors that may contribute to shaping the effects of drug pressure in a geographic region, similarities between the less genetically diverse and highly structured populations may facilitate the rapid spread and fixation of resistant alleles (Ariey et al. 2003; reviewed in Amato et al. 2018). Drug resistance mutations are often less fit than wildtype alleles (Hastings and Donnelly 2005), but reduced effective population size in low-transmission populations may allow these mutations to segregate (Escalante et al. 2009; Conrad et al. 2023) and subsequently fix by chance.

With multiple drug resistance alleles segregating in the *P. falciparum* populations, there is a need to understand the geographic location and time of origin of these mutations, as well as the identification of the targets of adaptation in the genome. We do not review the vast literature of drug resistance in *P. falciparum* here and instead refer the readers to previous reviews on the subject (Hastings and Donnelly 2005; Escalante et al. 2009; Takala-Harrison and Laufer 2015; Amato et al. 2018; Rosenthal et al. 2024). As new beneficial mutations increase in frequency and reach fixation, linked alleles hitchhike with the beneficial mutation to result in specific genomic signatures, referred to as a selective sweep. There is a decrease in nucleotide diversity near the beneficial fixation (Maynard Smith and Haigh 1974), a skew in the site frequency spectrum (Braverman et al. 1995), and a spatial pattern in linkage disequilibrium (Kim and Nielsen 2004), all of which are frequently used to detect loci where sweeps may have occurred (Nielsen 2005). While these methods have a reasonably high accuracy when selection is extremely strong and very recent, it becomes exceedingly difficult to detect sweeps that were only moderately strong or if the beneficial fixation occurred ∼0.5*N* generations ago (Przeworski 2002; Soni et al. 2023). It is especially difficult to distinguish between false and true positives if the population has experienced bottlenecks (Teshima et al. 2006; Crisci et al. 2013). Because within-host pathogen populations undergo strong recurrent bottlenecks every generation, it becomes important to account for nonequilibrium demography when attempting to identify loci involved in adaptation in natural populations. We consider some parameters that may specifically affect the detection of selective sweeps in *P. falciparum* populations below.

Because malaria populations drastically vary in their transmission dynamics across the globe, there is a large difference in the amount of effective recombination in different populations. For instance, African populations on average have much larger effective recombination rates than the South American populations, though transmission is highly heterogeneous at smaller geographic scales (see the section *Recombination*). This can result in very different lengths of genomic signatures due to sweeps (Figure 4). For instance, we estimate that in high-transmission populations, signatures of sweeps immediately post-fixation can extend to ∼6 kb when positive selection is weak (*s*=0.01) and up to 45 kb when selection is extremely strong (*s*=0.1). In contrast, in low-transmission populations, signatures of sweeps are expected to extend to much longer regions: ∼25 kb and 180 kb when selection is weak vs strong, respectively. Thus, while it might be easier to detect valleys of decreased diversity in low-transmission populations, which would span larger regions of the genome (up to 1/4th of the short chromosomes), high-transmission populations would allow for more precise identification of the target of selection due to smaller signatures around relevant loci, if selection was recent. These signatures of reduced diversity are significantly reduced if the sweep occurred *N*_e_ generations ago, which would be ∼9000-10,000 years ago, assuming that *P. falciparum* undergoes 8-9 generations (referring to all divisions in a transmission cycle) per year (reviewed in Ejigiri and Sinnis 2009; Smith et al. 2014; Vaughan and Kappe 2017; Graumans et al. 2020). Note that this is a lot more recent than the speciation event where *P. falciparum* jumped to the human host and started diverging from *P. prefalciparum* around 40,000-60,000 years ago (Otto, Gilabert, et al. 2018). Thus, while gene candidates involved in recently evolved drug resistance can be detected successfully, those involved in adaptation to the human host will be difficult to detect using standard population genetics approaches.

**Figure 4:**
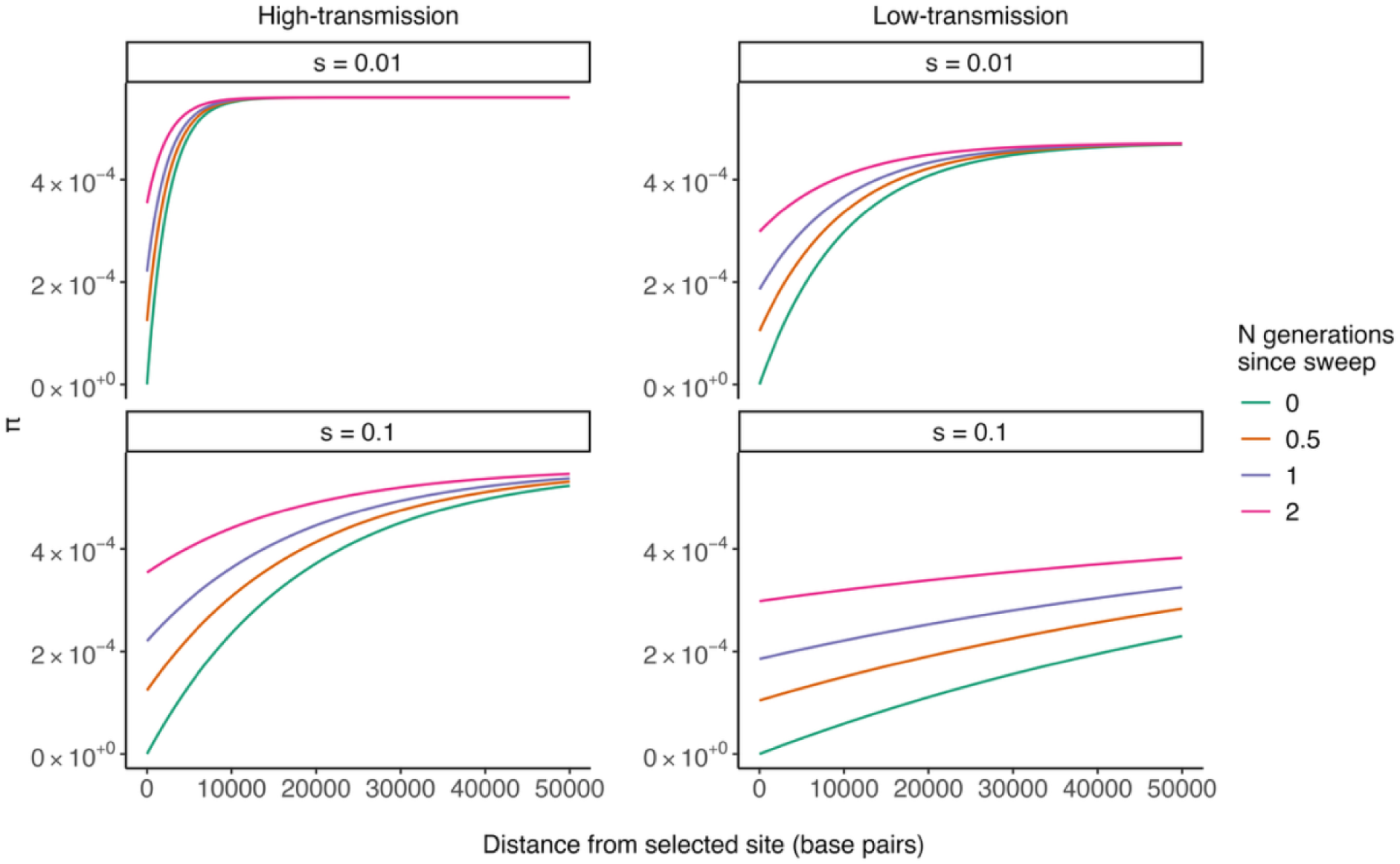
Expected signatures of selective sweeps in populations of high transmission (left) vs low transmission (right). Recovery of neutral nucleotide diversity (*π*) is shown as a function of the distance from the beneficial fixation. Here, we assume that the haploid Wright-Fisher population size (*N*) without accounting for inbreeding is 80,000 (estimated from (Chang et al. 2012; Otto et al. 2018), Wright’s inbreeding coefficient (*F*) was assumed to be 0.6 in the high-transmission and 0.9 in the low-transmission population. We assume that *N*_e_=*N*/(1+*F*), *r*_eff_ = *r*(1-*F*), and that the beneficial mutation is semidominant. The recovery of diversity post-fixation is calculated using equation 13 from Kim and Stephan (2000). Immediately post-fixation, neutral diversity recovers to 95% of its baseline value in ∼6 kb (*F*=0.6) and 25 kb (*F*=0.9) when selection is weaker (*s*=0.01) and in ∼46 kb (*F*=0.6), and 180 kb (*F*=0.9) when selection is extremely strong (*s*=0.1).

Consistent with variations in effective recombination rates across transmission dynamics, valleys of nucleotide diversity observed near genes responsible for drug-resistance have been reported to span much smaller regions in African populations: 3-10kb in Malawi (Ocholla et al. 2014), 10-13 kb in Ghana (Alam et al. 2011), 45kb in Kenya (Borrmann et al. 2013), and larger regions in populations of moderate to low transmission: 75-177kb at the China-Myanmar border (Wang et al. 2016), 600kb in Cambodia (Samad et al. 2015). It would be important to account for this difference in rates of recombination as well as time to origin for accurately estimating the selection coefficient of beneficial mutations in different populations.

Another important implication of vastly different recombination rates across populations is that the efficacy of selection, as well as the magnitude of indirect effects of selection, would be different across populations. Because low-transmission populations will have a reduced effective population size, beneficial mutations will be subjected to stronger genetic drift (*i.e.*, there will be more mutations with 2*N*_e_*s* < 1; Eyre-Walker and Keightley 2007). Secondly, populations with low transmission are likely to experience stronger linked effects of selection, more specifically, increased background selection as well as increased interference between selected alleles (Hill and Robertson 1966; Charlesworth and Jensen 2021). Both of these would decrease the probability of fixation of beneficial alleles in low-transmission populations. Moreover, as discussed above, the expected genomic size of signatures of recent selective sweeps would likely be highly population-specific. It might therefore be necessary to generate population-specific expectations to perform selection scans using the *P. falciparum* genome.

The malaria parasite has a unique life cycle (as discussed in the section *Infection Cycle*), such that the parasite population experiences subsequent rounds of expansions within a host, followed by bottlenecks during transmission to a new host. While the phases of population expansion result in stronger efficacy of purifying and positive selection, bottlenecks down to ∼10 parasites greatly reduce the effective population size, increasing the effects of genetic drift. Chang et al. (2013) evaluated the effect of the malaria life cycle on the effects of selection and found that compared to a Wright-Fisher (WF) population under equilibrium, the probability of fixation of beneficial mutations was drastically reduced, because new beneficial mutations are likely to be lost during infection bottlenecks. Moreover, the time to fixation of beneficial mutations was found to be larger in the malaria-specific demography, as opposed to a WF population under equilibrium. The longer time to fixation will allow for more time for linked neutral mutations to escape the haplotype carrying the beneficial mutation, resulting in less reduction of diversity due to sweeps compared to standard expectations. However, note that this study was restricted to understanding the dynamics of selected mutations *within* hosts. Thus, it is unclear how the effects of the within-host population size changes will affect evolutionary dynamics across the entire population.

Another important factor specific to the malaria life cycle is that meiosis occurs only once per generation (*i.e.*, the full life cycle) while the parasite replicates asexually (*i.e.*, without recombination) during the rest of the cycle. Interestingly, a recent simulation study (Ollivier et al. 2025) found that in organisms that undergo mitotic cell divisions interspersed infrequently with meiosis, hard sweeps can decrease diversity across all chromosomes. This is because during phases of clonal expansion, all chromosomes are fully linked to each other, extending hitchhiking effects genome-wide. This effect also lowers the baseline levels of neutral diversity genome-wide, in turn reducing the valley of diversity observed post-sweep. As *P. falciparum* undergoes a single meiosis event, which can be a result of selfing, every ∼35 mitotic divisions, a similar effect is likely to be observed, again reducing the expected effects of sweeps, possibly reducing the power to detect them. Thirdly, another factor pertinent to malaria populations is that the beneficial effects of mutations may be specific to particular stages of the life cycle. For instance, while drug resistance mutations have benefits within the human host, they are unlikely to be beneficial within the mosquito host, and in fact might even be costly (Segovia et al. 2025). A notable example of this is atovaquone, which selects for mutations in the parasite’s cytochrome *b* gene that contributes to drug resistance in humans, but are later lethal in the mosquito (Goodman et al. 2016). Similarly, mutations that allow better survival in the mosquito gut might play no role in increasing fitness in the human host. A systematic study about how such rapid temporal fluctuations in fitness will affect fixation probabilities and signatures of sweeps is currently lacking.

Finally, natural populations of the parasite have experienced a complex history with multiple migration and bottleneck events. *P. falciparum*, which is most closely related to the gorilla vector *P. praefalciparum*, diverged from their most recent common ancestor 40-60 thousand years ago (Otto, Gilabert, et al. 2018; Galaway et al. 2019). From Africa, the parasite spread globally, mirroring the migration of modern humans. Population structure observed today reflects both this ancient expansion and recent events, with African, South Asian, and South American parasites forming distinct, segregated genetic clusters. For example, Indian isolates segregate from Southeast Asian in PCA space, lying between Bangladesh/Myanmar and Thailand/Cambodia samples. Neighbor-joining trees support the clear separation between African and Asian lineages, and Indian samples occupy an intermediate position (Kumar et al. 2016). Meanwhile, South American isolates form a tight cluster closely related to African parasites, which may suggest the trans-Atlantic slave trade being the primary route of introduction (Lefebvre et al. 2023; Michel et al. 2024). Despite such complex demography, inference of selection coefficients in malaria populations has assumed a panmictic population at equilibrium (*e.g.*, Wootton et al. 2002; Nsanzabana et al. 2010), making these estimates less reliable. A careful study that incorporates the population history, including founder effects (e.g. in South American populations; (Lefebvre et al. 2023), along with the specifics of the malaria life cycle is needed to better estimate parameters of selection in malaria populations.

Recent studies have made available whole genome population genomic longitudinal datasets spanning pre-and-post intervention across populations of interest to quantify changes in allele and haplotype frequency in and near genes associated with drug resistance (Michel et al. 2024, Early et al. 2025, Cequeira et al. 2017, Amambua-Ngwa et al. 2026). Notably, community collaboration in the MalariaGEN Pf8 dataset (Abdel Hamid et al. 2025) has contributed to a more robust distribution of timepoints for whole-genome variant data across 34 countries, including Gambia (1966-2020), Kenya (1994-2020), and Thailand (2001-2018). Such datasets may enable more accurate inference of evolutionary parameters.

### TOWARDS BUILDING A BASELINE MODEL

An appropriate baseline model is essential to perform evolutionary inference from population genomic data and is especially challenging in pathogenic organisms due to their unique life cycles, complex demographic history with recurrent bottlenecks during transmission, strong bouts of selection against drugs, high-coding density across the genome, and skewed progeny distributions. Here, we have presented key considerations when building such a baseline model for inference in populations of the deadliest malaria parasite, *P. falciparum*. The complex evolutionary dynamics of *P. falciparum* require a baseline model that incorporates the effects of simultaneously acting evolutionary processes such as alternating cycles of asexual and sexual reproduction, selfing, transmission bottlenecks, and genome architecture (*i.e*., heterogeneity in coding density, recombination, and mutation rates). Importantly, an appropriate null model must adopt a framework that accounts for the within-host dynamics at shorter time scales and the global population history of multiple hosts over longer evolutionary times (Figure 5).

**Figure 5:**
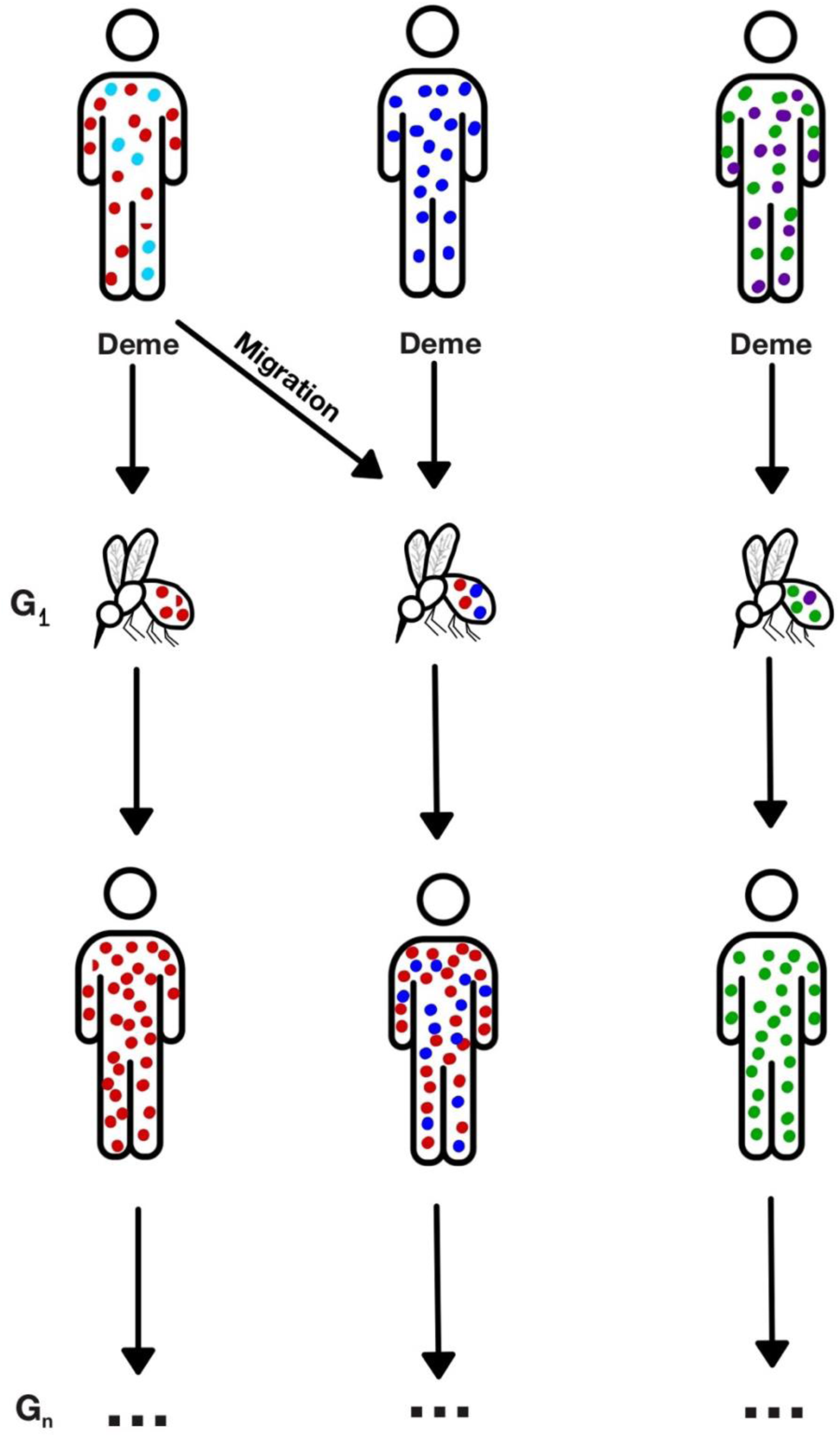
Depiction of a baseline evolutionary model for *P. falciparum* incorporating within-host dynamics (depicted by the solid circles that represent the parasite), between-host transmission (modeled as migration), and the long-term demographic history of the host metapopulation (where *G*_n_ represents the *n*^th^ generation).

This could be achieved by modeling a population of multiple hosts as a metapopulation composed of individual infected hosts as demes, with migration between demes representing superinfection. Population genetics has had a long tradition of considering a metapopulation framework to model structured populations (Wright 1943; Maruyama 1970). A metapopulation is a set of subpopulations or demes with some probability of exchanging migrants between them. We suggest that it may be possible to adopt a similar framework, albeit with a change, where each deme represents a human/mosquito host. Such an approach could allow for the flexibility to model the specifics of the life cycle, such as within-host clonal expansions and bottlenecks, as well as stochasticity during transmission between hosts (*i.e.*, allowing for extinctions of lineages).

These metapopulation models naturally bring up the idea of evolution at different time scales. There are two different time scales involved in the modeling of pathogen populations – evolution within hosts in the recent past and that across many hosts over many generations. Because individuals within a deme coalesce at a much shorter time scale than those across hosts, it is unclear whether evolution within hosts will affect the genealogy at longer time scales (Tellier and Lemaire 2014; Irwin et al. 2016). However, evolutionary processes of interest themselves occur at different time scales.

Dynamics of strongly beneficial mutations, like their fixation in the population, can occur at relatively short time scales. For instance, a new beneficial mutation in a haploid Wright-Fisher population takes on average 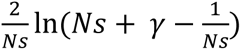 generations (scaled by *N*), conditional on fixation, where *N* is the effective population size, *s* is the selection coefficient, and gamma is Euler’s constant ∼0.577 (Hermisson and Pennings 2005). Thus, in a haploid population of effective size 50,000, it will take a selectively advantageous mutation (with *s*=0.01 and 0.1) ∼1,243 and ∼170 generations to reach fixation in the population, respectively. On the other hand, other evolutionarily important processes occur at longer timescales, which are of order *N* generations. For instance, in a haploid population, it takes on average *N* generations for a pair of genomes to coalesce, with a large standard deviation of *N* generations. Thus, patterns of variation present in the population sampled today are representative of the last 2*N* (100,000) generations in *P. falciparum* populations. This allows us to infer ancestral events such as ancient bottlenecks, migration out-of-Africa, and migration between countries that occurred in the past.

Processes related to the *P. falciparum* life cycle, like transmission bottlenecks, progeny skew due to a skewed offspring distribution during clonal expansion in the blood stage, and alternating sexual and asexual reproduction, are likely to have effects at shorter time scales. In particular, strong transmission bottlenecks can increase stochasticity and decrease the probability of fixation of beneficial mutations (Roze et al. 2005). Moreover, high variance in offspring distribution between strains can lead to non-WF dynamics, such as sweepstake reproduction within hosts, which increases the probability of fixation of beneficial mutations (Der et al. 2011). Studies suggest that the probability of fixation of beneficial mutations (Bessho and Otto 2022) and signatures of selective sweeps (Ollivier et al. 2025) are affected by alternating sexual and asexual reproduction. Thus, while characteristics of the life cycle are likely important in determining within-host evolution and how selected mutations behave in *P. falciparum* populations, it remains to be tested whether these processes affect long-term evolutionary outcomes of order *N* generations (see Box 1).

A process that can affect long-term evolutionary dynamics is the transmission of the parasite from one host to another. Highly stochastic and skewed transmission of malaria, where a few hosts may be more likely to infect other hosts, could lead to multiple mergers at longer time scales. Epidemiological studies suggest that mosquito biting is consistently over-dispersed across transmission settings with 20% of individuals responsible for 80% of transmission due to a combination of heterogeneous biting, as well as transmission bottlenecks (both at the human acquisition of infection and mosquito inoculation) (Graumans et al. 2020; Markwalter et al. 2024). This would lead to different expectations of neutral variant frequencies from those in a Wright-Fisher population (discussed in the section “Infection cycle”) and may be better explained by the Beta coalescent (Goldberg 2026). Again, note that if the number of hosts is large and the sampling is geographically scattered and species-wide, the long-term genealogy may converge to the Kingman coalescent (Wakeley and Aliacar 2001; Städler et al. 2009; Heuer and Sturm 2013). It is therefore not clear yet, which coalescent model would be the most appropriate null model for *P. falciparum* evolution, and it might depend on the scale of sampling.

Despite the lack of a known coalescent process, inference of the Ancestral Recombination Graph (ARG; McVean and Cardin 2005) could be insightful when inferring demographic history and population structure at the global level. However, because the rate of recombination in *P. falciparum* is ∼100-fold the rate of mutation, accurate construction of ARGs would be difficult using current methods. Alternatively, recent methods now allow for model-free inference of population size changes assuming the beta coalescent (Korfmann et al. 2024), which may be used if it can be established that that is the correct null for that population.

A large number of epidemiological models have been built to represent transmission in malaria populations (e.g., Watson et al. 2017; Whitlock et al. 2021). It may be possible to incorporate these parameters into population-genetic models. Moreover, it is becoming increasingly easier to perform agent-based forward simulations that incorporate mutation, recombination, selection, and demographic processes like recurrent bottlenecks (Haller and Messer). Such agent-based forward simulations that incorporate transmission dynamics learnt from epidemiological models may shed light on which coalescent model may be most appropriate for malaria populations and whether short-term processes should be accounted for when inferring processes at longer time scales. One such recent study (Hendry et al. 2021) found that while parasite prevalence mainly drives patterns of variation, different epidemiological variables that affect prevalence affect population genetics summary statistics differently. In particular, factors such as host clearance rate, biting rate, and the number of vectors determine the levels of statistics summarizing the SFS and LD differently. This study suggests that more complex models that bridge epidemiology and population genetics are required to model genetic variation in malaria populations. Similar simulation studies are needed to better understand whether the characteristics of the life cycle of *P. falciparum*, within-host evolution, and transmission dynamics between hosts affect short-term or long-term evolutionary processes (Box 1).

A major challenge for models that account for both within- and between-host evolution will be dealing with an extremely large parameter space and statistical identifiability. Epidemiological models already account for a large number of parameters (e.g., 10-45). A potential solution would be to obtain simplified theoretical models that summarize the effects of multiple parameters into effective population genetics quantities. For instance, it may be possible to adopt metapopulation models like the island model (Wright 1943) and the extinction-recolonization (Slatkin 1977) models to model pathogen populations, as they naturally allow a framework to incorporate within- and between-host evolution, as well as the effects of transmission bottlenecks. A combination of the Wright-Fisher and Moran models may be needed to account for overlapping generations. Further theoretical studies are required to understand whether an existing population genetics framework can be employed or whether new theoretical models are required to explain evolutionary dynamics in pathogen populations.

Alternatively, simulation studies can allow us to incorporate and vary many parameters simultaneously (as in Hendry et al. 2021). Additionally, simulations allow for incorporating uncertainty in evolutionary and biological parameters more easily by sampling from distributions rather than assuming constant values. This can be used to generate null expectations of patterns of variation and allow the development of accompanying simulation-based statistical approaches (*e*.*g*., Approximate Bayesian computation; Beaumont et al. 2002). For instance, we provide uncertainty around estimates of the parameters of the life cycle of *P. falciparum* in Supplementary Table 1 that may be utilized in simulations. On the other hand, simulations can be extremely time-consuming, especially if the number of parasites or the number of hosts being modeled is large, and if the parameter space being explored is large. Finally, in recombining populations like those of *P. falciparum*, summary statistics like nucleotide diversity are affected by long evolutionary times (∼10*N* generations), simulating which adds more computational time. With a recent explosion in new simulation tools, ranging from malaria-specific epidemiological simulations (Watson et al. 2017), to those combining malaria genetics and epidemiology (Hendry et al. 2021) and population genetics simulations for pathogen evolution in general (Xu et al. 2026), such endeavors can now be explored.

Finally, in order to account for the life cycle of *P. falciparum* and for both within- and between-host evolution simultaneously, a population genetics model that treats each haploid parasite genome individually is necessary. Because *P. falciparum* is a highly recombining eukaryotic species, and the mutation rate per site/cell division is similar to that of other unicellular eukaryotes, once the parasite population reaches 10^4^-10^5^ individuals within the host, there will be on average more than 58-575 new mutations in each parasite genome with many new mutations occurring on the same site in different individuals independently once the parasite population reaches 10^10^-10^13^ individuals. Although most new mutations will be rare, parasite diversity is unlikely to be captured accurately as a number of distinct strains (*e.g.,* multiplicity of infection). Consistent with this, deep sequencing studies of a single protein-coding gene within-host suggest hundreds of haplotypes (Juliano et al. 2010; Nkhoma et al. 2018). Moreover, if any of these new mutations were to generate a large phenotypic effect, then selection within-host will modify their allele frequency. Thus, a population genetics model that incorporates both within-host and between-host evolution will be necessary to describe all dynamics.

This recommendation conflicts with focusing on monoclonal or polyclonal samples as is currently common among malaria researchers and has been explicitly proposed as a theoretical model by Kwiatkowski (2024). Additionally, a challenge in implementing our recommendation is the lack of sequenced data that truly represents a single haploid genome. Multiple genomes of *P. falciparum* are typically condensed into monoclonal or polyclonal strains and collapsed into consensus genotypes, reducing the effective within-host diversity to a single haplotype. Additionally, many analyses remove polyclonal infections from analysis *(e.g.,* using thresholds such as *F_WS_* < 0.95). A recent study by Sabin et al. (2022) has shown that when using consensus sequences, one underestimates the level of diversity in the population, leading to an underestimation of the effective population size. Additionally, these data processing choices may bias the site frequency spectrum and distort patterns of linkage disequilibrium by removing recombinant haplotypes and minor clones, potentially biasing inference of demographic history and selection. Thus, while population genetics models may not explicitly keep track of the number of co-infection or super-infection events, these events would be a natural outcome of migration between and recombination within hosts, and it would be necessary for an appropriate model to accurately describe patterns of LD within and between hosts.

The ultimate goal for many researchers is to successfully detect targets of selection in the genome. While outlier-based methods will likely provide some candidates for targets of selection, it is unclear what proportion of loci experiencing recent selection will be present in the tails of the distribution of summary statistics. Besides asking how the life cycle and transmission dynamics affect neutral variation, it will be important moving forward to ask how they will affect the detection of targets of selection using outlier and model-based approaches (as done by Early et al. 2025). Implementing the correct null model will indeed be a difficult endeavour, but it will be essential for the accurate inference of population history, including patterns of global migration of the parasite, for predicting the evolution of drug resistance mutations in the malaria parasite, and to allow hypothesis testing, an essential component of basic science research.

## ACKNOWLEDGEMENTS

Research reported in this publication was supported by the National Institute of General Medical Sciences (R35GM154969 to PJ) and the National Institute of Allergy and Infectious Diseases (R01AI155730 and R21AI180675 to JL, R01AI189911 to JB), National Institutes of Health, as well as by Seed Funding provided to PJ by the College of Arts and Sciences, UNC, Chapel Hill. The funders had no role in study design, data collection and analysis, decision to publish, or preparation of the manuscript. We thank Jeff Jensen for thoroughly reading our manuscript and providing constructive comments that improved the manuscript. AI was not used in any part of the process of preparing this manuscript.

## Box 1.

**Key questions, challenges, and future directions in *P. falciparum* population genetics.** Potential topics of exploration may be categorized in a few major themes: A) Considerations to develop a realistic evolutionary null model that incorporates the parasite infection cycle and transmission dynamics, B) aspects to accurately quantify the evolutionary processes that underlie genetic diversity, and C) methods to improve approaches for inferring selection and demographic history in population genomic datasets.

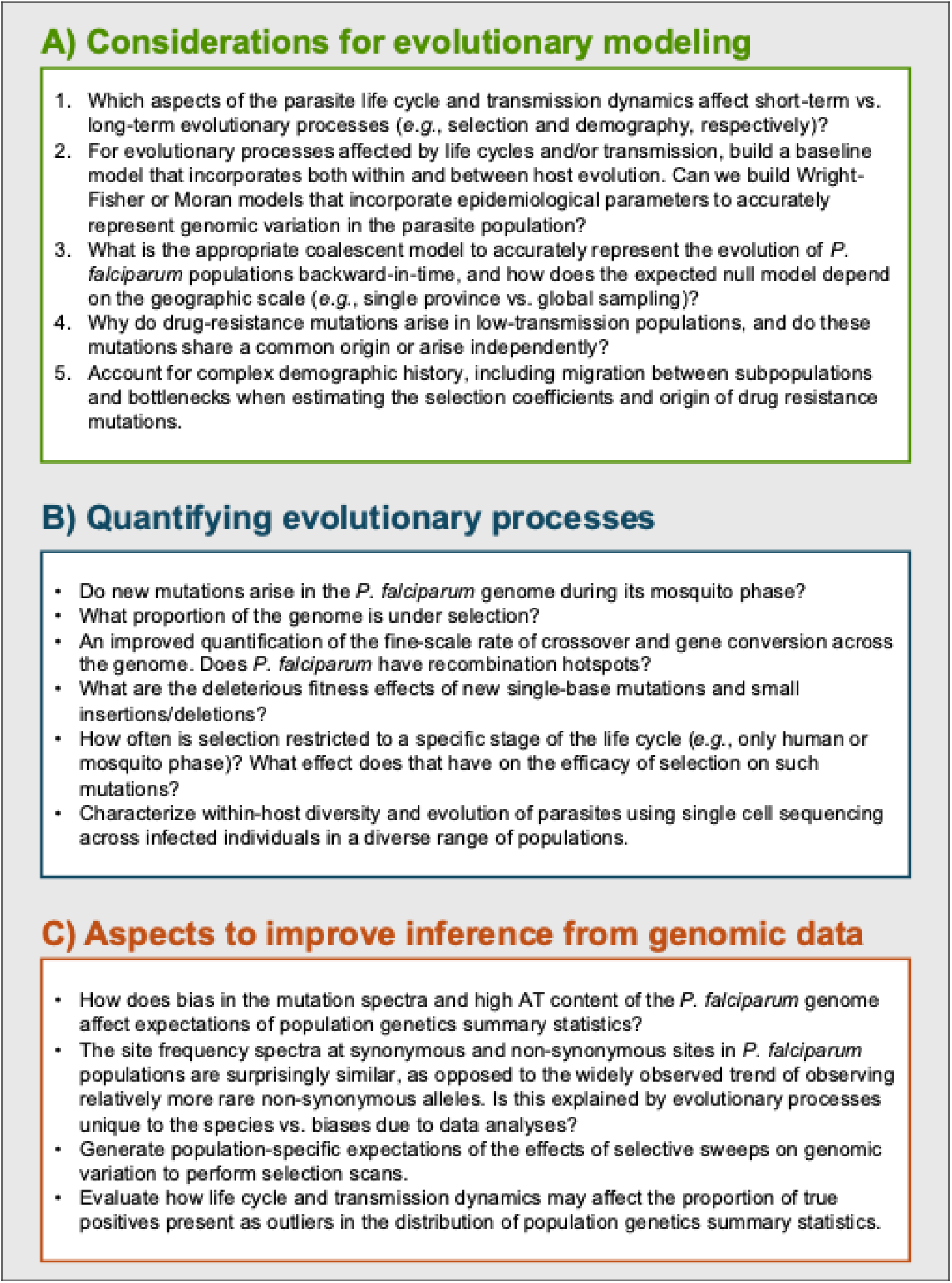

## SUPPLEMENTARY INFORMATION

**Table S1:** Previous estimates of the duration of parts of the lifecycle of *P. falciparum*, their approximate population sizes, and ploidy during the various stages of their life cycle. Variation in these estimates is indicated by the column “Overall range in the literature”.

**Table S2:** The proportion of time spent in each ploidy.

**Table S3.** Mean observed *P. falciparum F_WS_* values for all countries in the MalariaGEN Pf7 (2023).

**Table S4:** Theoretical expectations of neutral nucleotide diversity with BGS. Here we assume a Wright-Fisher population size of 80,000 haploid individuals and a mutation rate of 5.6 × 10^-9^ per site/ generation. We assumed *R_eff_* = *R*(1 − *F_it_*) where *F* = 0.6 for high-transmission and 0.9 for low-transmission populations, where *F* is the inbreeding coefficient. The genome-wide deleterious mutation rate (*U*) is obtained by assuming that 53% of the genome is under selection. When assuming a single selection coefficient, the relative diversity with BGS to that under neutrality, depicted by *B*, was calculated as *B*∼*exp*{−*U*/(2*s* + *R_self_*(1 − *s*))}. When assuming that selection coefficients follow a distribution of fitness effects (DFE), a combination of four non-overlapping uniform distributions was assumed, such that 0<*Ns*≤1, 1<*Ns*≤10, 10<*Ns*≤100, and 100 <*Ns*≤100,000, where *s* represents the disadvantage of a homozygous mutant relative to wild type. Each non-overlapping distribution was present in proportions *f*_0_, *f*_1_, *f*_2_, and *f*_3_, respectively. We assumed that *f*_0_=0, *f*_1_=0.333, *f*_2_=0.333, *f*_3_=0.333 at selected sites. When assuming a DFE, *B* was estimated using equations described in Marsh et al. (2025).

**Supplementary note A:** Estimation of the lower and upper bounds of the number of mitotic divisions during the human blood phase.

## REFERENCES

Abdel Hamid MM et al. 2023. Pf7: an open dataset of *Plasmodium falciparum* genome variation in 20,000 worldwide samples [version 1; peer review: 3 approved]. Wellcome Open Res. 8:22. 10.12688/wellcomeopenres.18681.1

Abdel Hamid MM et al. 2025. Pf8: an open dataset of *Plasmodium falciparum* genome variation in 33,325 worldwide samples [version 1; peer review: 3 approved]. Wellcome Open Res. 10:325. 10.12688/wellcomeopenres.24031.1

Achidi EA et al. 2008. A global network for investigating the genomic epidemiology of malaria. Nature. 456(7223):732–737. 10.1038/nature07632

Adam I et al. 2022. An open dataset of *Plasmodium vivax* genome variation in 1,895 worldwide samples. Wellcome Open Res. 7:136. 10.12688/wellcomeopenres.17795.1

Ahouidi A et al. 2021. An open dataset of *Plasmodium falciparum* genome variation in 7,000 worldwide samples [version 2; peer review: 2 approved]. Wellcome Open Res. 6:42. 10.12688/wellcomeopenres.16168.2

Alam MT et al. 2011. Selective sweeps and genetic lineages of *Plasmodium falciparum* drug-resistant alleles in Ghana. J Infect Dis. 203(2):220–227. 10.1093/infdis/jiq038

Amambua-Ngwa A et al. 2019. Major subpopulations of *Plasmodium falciparum* in sub-Saharan Africa. Science. 365(6455):813–816. 10.1126/science.aav5427

Amaratunga C et al. 2016. Dihydroartemisinin-piperaquine resistance in *Plasmodium falciparum* malaria in Cambodia: a multisite prospective cohort study. Lancet Infect Dis. 16(3):357–365. 10.1016/S1473-3099(15)00487-9

Amato R et al. 2018. Origins of the current outbreak of multidrug-resistant malaria in Southeast Asia: retrospective genetic study. Lancet Infect Dis. 18(3):337–345. 10.1016/S1473-3099(18)30068-9

Amino R et al. 2006. Quantitative imaging of *Plasmodium* transmission from mosquito to mammal. Nat Med. 12(2):220–224. 10.1038/nm1350

Anderson TJC et al. 2000. Microsatellite markers reveal a spectrum of population structures in the malaria parasite *Plasmodium falciparum*. Mol Biol Evol. 17(10):1467–1482. 10.1093/oxfordjournals.molbev.a026247

Andolina C et al. 2024. A transmission bottleneck for malaria? Quantification of sporozoite expelling by *Anopheles* mosquitoes infected with laboratory and naturally circulating *P. falciparum* gametocytes. eLife. 12:RP90989. 10.7554/eLife.90989.2

Ariey F, Duchemin J-B, Robert V. 2003. Metapopulation concepts applied to falciparum malaria and their impacts on the emergence and spread of chloroquine resistance. Infect Genet Evol. 2(3):185–192. 10.1016/S1567-1348(02)00099-0

Arrow KJ, Panosian C, Gelband H. 2004a. A Brief History of Malaria. In: Saving Lives, Buying Time: Economics of Malaria Drugs in an Age of Resistance. National Academies Press (US) https://www.ncbi.nlm.nih.gov/books/NBK215638/

Arrow KJ, Panosian C, Gelband H. 2004b. The Parasite, the Mosquito, and the Disease. In: Saving Lives, Buying Time: Economics of Malaria Drugs in an Age of Resistance. National Academies Press (US) https://www.ncbi.nlm.nih.gov/books/NBK215619/

Auburn S et al. 2012. Characterization of within-host *Plasmodium falciparum* diversity using next-generation sequence data. PLoS One. 7(2):e32891. 10.1371/journal.pone.0032891

Auton A et al. 2015. A global reference for human genetic variation. Nature. 526(7571):68–74. 10.1038/nature15393

Bacon DJ et al. 2009. Dynamics of malaria drug resistance patterns in the Amazon basin region following changes in Peruvian national treatment policy for uncomplicated malaria. Antimicrob Agents Chemother. 53(5):2042–2051. 10.1128/aac.01677-08

Bank C et al. 2014. Thinking too positive? Revisiting current methods of population genetic selection inference. Trends Genet. 30(12):540–546. 10.1016/j.tig.2014.09.010

Basu L et al. 2025. Drug resistance and new strategies of prevention against malaria: an ongoing battle. J Vector Borne Dis. 62(1):9–15. 10.4103/JVBD.JVBD_72_24

Baton LA, Ranford-Cartwright LC. 2005. Spreading the seeds of million-murdering death: metamorphoses of malaria in the mosquito. Trends Parasitol. 21(12):573–580. 10.1016/j.pt.2005.09.012

Baudat F et al. 2010. PRDM9 is a major determinant of meiotic recombination hotspots in humans and mice. Science. 327(5967):836–840. 10.1126/science.1183439

Beaumont MA, Zhang W, Balding DJ. 2002. Approximate Bayesian computation in population genetics. Genetics. 162(4):2025–2035. 10.1093/genetics/162.4.2025

Begun DJ, Aquadro CF. 1992. Levels of naturally occurring DNA polymorphism correlate with recombination rates in *D. melanogaster*. Nature. 356(6369):519–520. 10.1038/356519a0

Behrens HM, Spielmann T. 2024. Identification of domains in *Plasmodium falciparum* proteins of unknown function using DALI search on AlphaFold predictions. Sci Rep. 14(1):10527. 10.1038/s41598-024-60058-x

Bessho K, Otto SP. 2022. Fixation and effective size in a haploid–diploid population with asexual reproduction. Theor Popul Biol. 143:30–45. 10.1016/j.tpb.2021.11.002

Birkner M et al. 2009. A modified lookdown construction for the Xi-Fleming-Viot process with mutation and populations with recurrent bottlenecks. Alea Lat Am J Probab Math Stat. 6:25–61. 10.48550/ARXIV.0808.0412

Blath J, Cronjäger MC, Eldon B, Hammer M. 2016. The site-frequency spectrum associated with Ξ-coalescents. Theor Popul Biol. 110:36–50. 10.1016/j.tpb.2016.04.002

Bopp SER et al. 2013. Mitotic evolution of *Plasmodium falciparum* shows a stable core genome but recombination in antigen families. PLoS Genet. 9(2):e1003293. 10.1371/journal.pgen.1003293

Borrmann S et al. 2013. Genome-wide screen identifies new candidate genes associated with artemisinin susceptibility in *Plasmodium falciparum* in Kenya. Sci Rep. 3(1):3318. 10.1038/srep03318

Brackney DE, LaReau JC, Smith RC. 2021. Frequency matters: how successive feeding episodes by blood-feeding insect vectors influences disease transmission. PLoS Pathog. 17(6):e1009590. 10.1371/journal.ppat.1009590

Brandvain Y, Wright SI. 2016. The limits of natural selection in a nonequilibrium world. Trends Genet. 32(4):201–210. 10.1016/j.tig.2016.01.004

Braverman JM et al. 1995. The hitchhiking effect on the site frequency spectrum of DNA polymorphisms. Genetics. 140(2):783–796. 10.1093/genetics/140.2.783

Brhane BG et al. 2025. Rising prevalence of *Plasmodium falciparum* artemisinin resistance mutations in Ethiopia. Commun Med. 5(1):297. 10.1038/s43856-025-01008-0

Briggs J et al. 2020. Sex-based differences in clearance of chronic *Plasmodium falciparum* infection. eLife. 9:e59872. 10.7554/eLife.59872

Burger G, Moreira S, Valach M. 2016. Genes in hiding. Trends Genet. 32(9):553–565. 10.1016/j.tig.2016.06.005

Button-Simons KA et al. 2021. The power and promise of genetic mapping from *Plasmodium falciparum* crosses utilizing human liver-chimeric mice. Commun Biol. 4(1):734. 10.1038/s42003-021-02210-1

Cai H et al. 2012. Module-based subnetwork alignments reveal novel transcriptional regulators in malaria parasite *Plasmodium falciparum*. BMC Syst Biol. 6(Suppl 3):S5. 10.1186/1752-0509-6-S3-S5

Camponovo F, Buckee CO, Taylor AR. 2023. Measurably recombining malaria parasites. Trends Parasitol. 39(1):17–25. 10.1016/j.pt.2022.11.002

Carey-Ewend K et al. 2024. Population genomics of *Plasmodium ovale* species in sub-Saharan Africa. Nat Commun. 15(1):10297. 10.1038/s41467-024-54667-3

Carlson CS et al. 2005. Genomic regions exhibiting positive selection identified from dense genotype data. Genome Res. 15(11):1553–1565. 10.1101/gr.4326505

Carlton JM et al. 2008. Comparative genomics of the neglected human malaria parasite *Plasmodium vivax*. Nature. 455(7214):757–763. 10.1038/nature07327

Carrasquilla M et al. 2022. Resolving drug selection and migration in an inbred South American *Plasmodium falciparum* population with identity-by-descent analysis. PLoS Pathog. 18(12):e1010993. 10.1371/journal.ppat.1010993

Carter LM et al. 2013. Stress and sex in malaria parasites. Evol Med Public Health. 2013(1):135–147. 10.1093/emph/eot011

CDC. 2024. Malaria; [accessed 2025 Aug 15]. https://www.cdc.gov/malaria/php/impact/index.html

Chan S, Ch’ng J-H, Wahlgren M, Thutkawkorapin J. 2017. Frequent GU wobble pairings reduce translation efficiency in *Plasmodium falciparum*. Sci Rep. 7(1):723. 10.1038/s41598-017-00801-9

Chang H-H et al. 2012. Genomic sequencing of *Plasmodium falciparum* malaria parasites from Senegal reveals the demographic history of the population. Mol Biol Evol. 29(11):3427–3439. 10.1093/molbev/mss161

Chang H-H et al. 2013. Malaria life cycle intensifies both natural selection and random genetic drift. Proc Natl Acad Sci USA. 110(50):20129–20134. 10.1073/pnas.1319857110

Charlesworth B. 2012. The effects of deleterious mutations on evolution at linked sites. Genetics. 190(1):5–22. 10.1534/genetics.111.134288

Charlesworth B. 2013. Background Selection 20 Years on: The Wilhelmine E. Key 2012 Invitational Lecture. J Hered. 104(2):161–171. 10.1093/jhered/ess136

Charlesworth B, Jensen JD. 2021. Effects of selection at linked sites on patterns of genetic variability. Annu Rev Ecol Evol Syst. 52(1):177–197. 10.1146/annurev-ecolsys-010621-044528

Charlesworth B, Morgan MT, Charlesworth D. 1993. The effect of deleterious mutations on neutral molecular variation. Genetics. 134(4):1289–1303. 10.1093/genetics/134.4.1289

Chavasse D. 2002. Know your enemy. Malawi Med J. 14(1):7–8

Chawla J, Oberstaller J, Adams JH. 2021. Targeting gametocytes of the malaria parasite *Plasmodium falciparum* in a functional genomics era: next steps. Pathogens. 10(3):346. 10.3390/pathogens10030346

Chenet SM et al. 2015. Longitudinal analysis of *Plasmodium falciparum* genetic variation in Turbo, Colombia: implications for malaria control and elimination. Malar J. 14(1):363. 10.1186/s12936-015-0887-9

Churcher TS et al. 2012. Measuring the blockade of malaria transmission – an analysis of the standard membrane feeding assay. Int J Parasitol. 42(11):1037–1044. 10.1016/j.ijpara.2012.09.002

Claessens A et al. 2014. Generation of antigenic diversity in *Plasmodium falciparum* by structured rearrangement of *Var* genes during mitosis. PLoS Genet. 10(12):e1004812. 10.1371/journal.pgen.1004812

Collins KA et al. 2018. A controlled human malaria infection model enabling evaluation of transmission-blocking interventions. J Clin Invest. 128(4):1551–1562. 10.1172/JCI98012

Comeron JM, Ratnappan R, Bailin S. 2012. The many landscapes of recombination in *Drosophila melanogaster*. PLoS Genet. 8(10):e1002905. 10.1371/journal.pgen.1002905

Conrad MD et al. 2023. Evolution of partial resistance to artemisinins in malaria parasites in Uganda. N Engl J Med. 389(8):722–732. 10.1056/NEJMoa2211803

Cowman AF et al. 1988. Amino acid changes linked to pyrimethamine resistance in the dihydrofolate reductase-thymidylate synthase gene of *Plasmodium falciparum*. Proc Natl Acad Sci USA. 85(23):9109–9113. 10.1073/pnas.85.23.9109

Cowman AF, Crabb BS. 2006. Invasion of red blood cells by malaria parasites. Cell. 124(4):755–766. 10.1016/j.cell.2006.02.006

Crellen T, Iantorno S. 2015. A switch in time. Nat Rev Microbiol. 13(4):190–190. 10.1038/nrmicro3458

Crisci JL, Poh Y-P, Mahajan S, Jensen JD. 2013. The impact of equilibrium assumptions on tests of selection. Front Genet. 4. 10.3389/fgene.2013.00235

Cutter AD, Payseur BA. 2013. Genomic signatures of selection at linked sites: unifying the disparity among species. Nat Rev Genet. 14(4):262–274. 10.1038/nrg3425

Davydov EV et al. 2010. Identifying a high fraction of the human genome to be under selective constraint using GERP++. PLoS Comput Biol. 6(12):e1001025. 10.1371/journal.pcbi.1001025

Der R, Epstein CL, Plotkin JB. 2011. Generalized population models and the nature of genetic drift. Theor Popul Biol. 80(2):80–99. 10.1016/j.tpb.2011.06.004

Doumbe-Belisse P et al. 2018. High malaria transmission sustained by *Anopheles gambiae s.l.* occurring both indoors and outdoors in the city of Yaoundé, Cameroon. Wellcome Open Res. 3:164. 10.12688/wellcomeopenres.14963.1

Duchemin J-B et al. 2003. Malaria transmission in urban sub-Saharan Africa. Am J Trop Med Hyg. 68(2):169–176. 10.4269/ajtmh.2003.68.169

Early AM, Pelleau S, Musset L, Neafsey DE. 2025. Temporal patterns of haplotypic and allelic diversity reflect the changing selection landscape of the malaria parasite *Plasmodium falciparum* Rogers R, editor. Mol Biol Evol. 42(4):msaf075. 10.1093/molbev/msaf075

Echeverry DF et al. 2013. Long term persistence of clonal malaria parasite *Plasmodium falciparum* lineages in the Colombian pacific region. BMC Genet. 14(1):2. 10.1186/1471-2156-14-2

Ejigiri I, Sinnis P. 2009. *Plasmodium* sporozoite-host interactions from the dermis to the hepatocyte. Curr Opin Microbiol. 12(4):401–407. 10.1016/j.mib.2009.06.006

Eldon B, Wakeley J. 2006. Coalescent processes when the distribution of offspring number among individuals is highly skewed. Genetics. 172(4):2621–2633. 10.1534/genetics.105.052175

Eldon B, Wakeley J. 2008. Linkage disequilibrium under skewed offspring distribution among individuals in a population. Genetics. 178(3):1517–1532. 10.1534/genetics.107.075200

Eldon B, Wakeley J. 2009. Coalescence times and FST under a skewed offspring distribution among individuals in a population. Genetics. 181(2):615–629. 10.1534/genetics.108.094342

Escalante AA, Smith DL, Kim Y. 2009. The dynamics of mutations associated with anti-malarial drug resistance in *Plasmodium falciparum*. Trends Parasitol. 25(12):557–563. 10.1016/j.pt.2009.09.008

Ewing GB, Jensen JD. 2016. The consequences of not accounting for background selection in demographic inference. Mol Ecol. 25(1):135–141. 10.1111/mec.13390

Eyre-Walker A, Keightley PD. 2007. The distribution of fitness effects of new mutations. Nat Rev Genet. 8(8):610–618. 10.1038/nrg2146

Frosch AEP et al. 2014. Return of widespread chloroquine-sensitive *Plasmodium falciparum* to Malawi. J Infect Dis. 210(7):1110–1114. 10.1093/infdis/jiu216

Galaway F et al. 2019. Resurrection of the ancestral RH5 invasion ligand provides a molecular explanation for the origin of *P. falciparum* malaria in humans. PLoS Biol. 17(10):e3000490. 10.1371/journal.pbio.3000490

Gao Z, Wyman MJ, Sella G, Przeworski M. 2016. Interpreting the dependence of mutation rates on age and time. PLoS Biol. 14(1):e1002355. 10.1371/journal.pbio.1002355

Gardner KB et al. 2011. Protein-based signatures of functional evolution in *Plasmodium falciparum*. BMC Evol Biol. 11(1):257. 10.1186/1471-2148-11-257

Gardner MJ et al. 1998. Chromosome 2 sequence of the human malaria parasite *Plasmodium falciparum*. Science. 282(5391):1126–1132. 10.1126/science.282.5391.1126

Gardner MJ et al. 2002. Genome sequence of the human malaria parasite *Plasmodium falciparum*. Nature. 419(6906):498–511. 10.1038/nature01097

Garud NR, Messer PW, Buzbas EO, Petrov DA. 2015. Recent selective sweeps in North American *Drosophila melanogaster* show signatures of soft sweeps. PLoS Genet. 11(2):e1005004. 10.1371/journal.pgen.1005004

Goldberg A. 2026. Rare variation in malaria parasites biases population-genetic inference. 10.64898/2026.01.09.698000

Goodman CD et al. 2016. Parasites resistant to the antimalarial atovaquone fail to transmit by mosquitoes. Science. 352(6283):349–353. 10.1126/science.aad9279

Gouagna LC et al. 1998. The early sporogonic cycle of *Plasmodium falciparum* in laboratory-infected *Anopheles gambiae*: an estimation of parasite efficacy. Trop Med Int Health. 3(1):21–28. 10.1046/j.1365-3156.1998.00156.x

Graumans W, Jacobs E, Bousema T, Sinnis P. 2020. When is a *Plasmodium*-infected mosquito an infectious mosquito? Trends Parasitol. 36(8):705–716. 10.1016/j.pt.2020.05.011

Griffing S et al. 2010. *Pfmdr1* amplification and fixation of *pfcrt* chloroquine resistance alleles in *Plasmodium falciparum* in Venezuela. Antimicrob Agents Chemother. 54(4):1572–1579. 10.1128/AAC.01243-09

Guelbéogo WM et al. 2018. Variation in natural exposure to *Anopheles* mosquitoes and its effects on malaria transmission. eLife. 7:e32625. 10.7554/eLife.32625

Guo B et al. 2024. Strong positive selection biases identity-by-descent-based inferences of recent demography and population structure in *Plasmodium falciparum*. Nat Commun. 15(1):2499. 10.1038/s41467-024-46659-0

Hagenah LM et al. 2024. Additional PfCRT mutations driven by selective pressure for improved fitness can result in the loss of piperaquine resistance and altered *Plasmodium falciparum* physiology. mBio. 15(1):e01832–23. 10.1128/mbio.01832-23

Hailemeskel E et al. 2024. Dynamics of asymptomatic *Plasmodium falciparum* and *Plasmodium vivax* infections and their infectiousness to mosquitoes in a low transmission setting of Ethiopia: a longitudinal observational study. Int J Infect Dis. 143:107010. 10.1016/j.ijid.2024.107010

Haller BC, Messer PW. SLiM: An Evolutionary Simulation Framework.

Halligan DL, Keightley PD. 2009. Spontaneous mutation accumulation studies in evolutionary genetics. Annu Rev Ecol Evol Syst. 40(1):151–172. 10.1146/annurev.ecolsys.39.110707.173437

Hamilton WL et al. 2017. Extreme mutation bias and high AT content in *Plasmodium falciparum*. Nucleic Acids Res. 45(4):1889–1901. 10.1093/nar/gkw1259

Hastings IM, Donnelly MJ. 2005. The impact of antimalarial drug resistance mutations on parasite fitness, and its implications for the evolution of resistance. Drug Resist Updat. 8(1):43–50. 10.1016/j.drup.2005.03.003

Hawking F, Worms MJ, Gammage K. 1968. 24- and 48-hour cycles of malaria parasites in the blood; their purpose, production and control. Trans R Soc Trop Med Hyg. 62(6):731–760. 10.1016/0035-9203(68)90001-1

Hayton K et al. 2008. Erythrocyte binding protein PfRH5 polymorphisms determine species-specific pathways of *Plasmodium falciparum* invasion. Cell Host Microbe. 4(1):40–51. 10.1016/j.chom.2008.06.001

Hendry JA, Kwiatkowski D, McVean G. 2021. Elucidating relationships between *P. falciparum* prevalence and measures of genetic diversity with a combined genetic-epidemiological model of malaria. PLoS Comput Biol. 17(8):e1009287. 10.1371/journal.pcbi.1009287

Hermisson J, Pennings PS. 2005. Soft sweeps. Genetics. 169(4):2335–2352. 10.1534/genetics.104.036947

Heuer B, Sturm A. 2013. On spatial coalescents with multiple mergers in two dimensions. Theor Popul Biol. 87:90–104. 10.1016/j.tpb.2012.11.006

Hill WG, Babiker HA, Ranford-Cartwright LC, Walliker D. 1995. Estimation of inbreeding coefficients from genotypic data on multiple alleles, and application to estimation of clonality in malaria parasites. Genet Res. 65(1):53–61. 10.1017/S0016672300033000

Hill WG, Robertson A. 1966. The effect of linkage on limits to artificial selection. Genet Res. 8(3):269–294. 10.1017/S0016672300010156

Holzschuh A et al. 2024. *Plasmodium falciparum* transmission in the highlands of Ethiopia is driven by closely related and clonal parasites. Mol Ecol. 33(6):e17292. 10.1111/mec.17292

Imwong M et al. 2017. Spread of a single multidrug resistant malaria parasite lineage (PfPailin) to Vietnam. Lancet Infect Dis. 17(10):1022–1023. 10.1016/S1473-3099(17)30524-8

Irwin KK et al. 2016. On the importance of skewed offspring distributions and background selection in virus population genetics. Heredity. 117(6):393–399. 10.1038/hdy.2016.58

Jeffares DC et al. 2007. Genome variation and evolution of the malaria parasite *Plasmodium falciparum*. Nat Genet. 39(1):120–125. 10.1038/ng1931

Jeffreys AJ, May CA. 2004. Intense and highly localized gene conversion activity in human meiotic crossover hot spots. Nat Genet. 36(2):151–156. 10.1038/ng1287

Jensen-Seaman MI et al. 2004. Comparative recombination rates in the rat, mouse, and human genomes. Genome Res. 14(4):528–538. 10.1101/gr.1970304

Jiang H et al. 2011. High recombination rates and hotspots in a *Plasmodium falciparum* genetic cross. Genome Biol. 12(4):R33. 10.1186/gb-2011-12-4-r33

Johri P, Riall K, et al. 2021. The impact of purifying and background selection on the inference of population history: problems and prospects. Mol Biol Evol. 38(7):2986–3003. 10.1093/molbev/msab050

Johri P, Charlesworth B, et al. 2021. Revisiting the notion of deleterious sweeps. Genetics. 219(3):iyab094. 10.1093/genetics/iyab094

Johri P, Aquadro CF, et al. 2022. Recommendations for improving statistical inference in population genomics. PLoS Biol. 20(5):e3001669. 10.1371/journal.pbio.3001669

Johri P, Charlesworth B, Jensen JD. 2020. Toward an evolutionarily appropriate null model: jointly inferring demography and purifying selection. Genetics. 215(1):173–192. 10.1534/genetics.119.303002

Johri P, Marinov GK, Doak TG, Lynch M. 2019. Population genetics of *Paramecium* mitochondrial genomes: recombination, mutation spectrum, and efficacy of selection. Genome Biol Evol. 11(5):1398–1416. 10.1093/gbe/evz081

Johri P, Pfeifer SP, Jensen JD. 2023. Developing an evolutionary baseline model for humans: jointly inferring purifying selection with population history. Mol Biol Evol. 40(5):msad100. 10.1093/molbev/msad100

Johri P, Stephan W, Jensen JD. 2022. Soft selective sweeps: addressing new definitions, evaluating competing models, and interpreting empirical outliers. PLoS Genet. 18(2):e1010022. 10.1371/journal.pgen.1010022

Juliano JJ et al. 2010. Exposing malaria in-host diversity and estimating population diversity by capture-recapture using massively parallel pyrosequencing. Proc Natl Acad Sci USA. 107(46):20138–20143. 10.1073/pnas.1007068107

Kanatani S et al. 2024. Revisiting the *Plasmodium* sporozoite inoculum and elucidating the efficiency with which malaria parasites progress through the mosquito. Nat Commun. 15(1):748. 10.1038/s41467-024-44962-4

Kappe SHI, Vaughan AM, Boddey JA, Cowman AF. 2010. That was then but this is now: malaria research in the time of an eradication agenda. Science. 328(5980):862–866. 10.1126/science.1184785

Kelley JL et al. 2006. Genomic signatures of positive selection in humans and the limits of outlier approaches. Genome Res. 16(8):980–989. 10.1101/gr.5157306

Kerr PJ, Ranford-Cartwright LC, Walliker D. 1994. Proof of intragenic recombination in *Plasmodium falciparum*. Mol Biochem Parasitol. 66(2):241–248. 10.1016/0166-6851(94)90151-1

Kibota TT, Lynch M. 1996. Estimate of the genomic mutation rate deleterious to overall fitness in *E. coli*. Nature. 381(6584):694–696. 10.1038/381694a0

Kim Y, Nielsen R. 2004. Linkage disequilibrium as a signature of selective sweeps. Genetics. 167(3):1513–1524. 10.1534/genetics.103.025387

Kim Y, Stephan W. 2000. Joint effects of genetic hitchhiking and background selection on neutral variation. Genetics. 155(3):1415–1427. 10.1093/genetics/155.3.1415

Kingman JFC. 1982. The coalescent. Stoch Process Their Appl. 13(3):235–248. 10.1016/0304-4149(82)90011-4

Korfmann K et al. 2024. Simultaneous inference of past demography and selection from the ancestral recombination graph under the beta coalescent. Peer Community J. 4:e33. 10.24072/pcjournal.397

Kublin JG et al. 2003. Reemergence of chloroquine-sensitive *Plasmodium falciparum* malaria after cessation of chloroquine use in Malawi. J Infect Dis. 187(12):1870–1875. 10.1086/375419

Kumar S et al. 2016. Distinct genomic architecture of *Plasmodium falciparum* populations from South Asia. Mol Biochem Parasitol. 210(1):1–4. 10.1016/j.molbiopara.2016.07.005

Kwiatkowski D. 2024. Modelling transmission dynamics and genomic diversity in a recombining parasite population. Wellcome Open Res. 9:215. 10.12688/wellcomeopenres.19092.1

Lambert B, North A, Godfray HCJ. 2022. A meta-analysis of longevity estimates of mosquito vectors of disease. 2022.05.30.494059. 10.1101/2022.05.30.494059

Lefebvre MJM et al. 2023. Population genomic evidence of adaptive response during the invasion history of *Plasmodium falciparum* in the Americas. Mol Biol Evol. 40(5):msad082. 10.1093/molbev/msad082

Li X et al. 2019. Genetic mapping of fitness determinants across the malaria parasite *Plasmodium falciparum* life cycle. PLoS Genet. 15(10):e1008453. 10.1371/journal.pgen.1008453

Lin JT, Saunders DL, Meshnick SR. 2014. The role of submicroscopic parasitemia in malaria transmission: what is the evidence? Trends Parasitol. 30(4):183–190. 10.1016/j.pt.2014.02.004

Lindblade KA et al. 2013. The silent threat: asymptomatic parasitemia and malaria transmission. Expert Rev Anti-Infect Ther. 11(6):623–639. 10.1586/eri.13.45

Lovett ST. 2004. Encoded errors: Mutations and rearrangements mediated by misalignment at repetitive DNA sequences. Mol Microbiol. 52(5):1243–1253. 10.1111/j.1365-2958.2004.04076.x

Lynch M. 2006. The origins of eukaryotic gene structure. Mol Biol Evol. 23(2):450–468. 10.1093/molbev/msj050

Lynch M et al. 2016. Genetic drift, selection and the evolution of the mutation rate. Nat Rev Genet. 17(11):704–714. 10.1038/nrg.2016.104

Lynch M, Conery JS. 2003. The origins of genome complexity. Science. 302(5649):1401–1404. 10.1126/science.1089370

Mackay TFC et al. 2012. The *Drosophila melanogaster* Genetic Reference Panel. Nature. 482(7384):173–178. 10.1038/nature10811

Mancera E et al. 2008. High-resolution mapping of meiotic crossovers and non-crossovers in yeast. Nature. 454(7203):479–485. 10.1038/nature07135

Manske M et al. 2012. Analysis of *Plasmodium falciparum* diversity in natural infections by deep sequencing. Nature. 487(7407):375–379. 10.1038/nature11174

Markwalter CF et al. 2024. *Plasmodium falciparum* infection in humans and mosquitoes influence natural Anopheline biting behavior and transmission. Nat Commun. 15(1):4626. 10.1038/s41467-024-49080-9

Maruyama T. 1970. Effective number of alleles in a subdivided population. Theor Popul Biol. 1(3):273–306. 10.1016/0040-5809(70)90047-X

Maynard Smith J, Haigh J. 1974. The hitch-hiking effect of a favourable gene. Genet Res. 23(1):23–35

McDew-White M et al. 2019. Mode and tempo of microsatellite length change in a malaria parasite mutation accumulation experiment. Genome Biol Evol. 11(7):1971–1985. 10.1093/gbe/evz140

McVean GAT, Cardin NJ. 2005. Approximating the coalescent with recombination. Phil Trans R Soc B. 360(1459):1387–1393. 10.1098/rstb.2005.1673

Michel M et al. 2024. Ancient *Plasmodium* genomes shed light on the history of human malaria. Nature. 631(8019):125–133. 10.1038/s41586-024-07546-2

Miles A et al. 2016. Indels, structural variation, and recombination drive genomic diversity in *Plasmodium falciparum*. Genome Res. 26(9):1288–1299. 10.1101/gr.203711.115

Miotto O et al. 2013. Multiple populations of artemisinin-resistant *Plasmodium falciparum* in Cambodia. Nat Genet. 45(6):648–655. 10.1038/ng.2624

Mita T et al. 2007. Independent evolution of pyrimethamine resistance in *Plasmodium falciparum* isolates in Melanesia. Antimicrob Agents Chemother. 51(3):1071–1077. 10.1128/aac.01186-06

Mita T et al. 2011. Limited geographical origin and global spread of sulfadoxine-resistant *dhps* alleles in *Plasmodium falciparum* populations. J Infect Dis. 204(12):1980–1988. 10.1093/infdis/jir664

Möhle M, Sagitov S. 2001. A classification of coalescent processes for haploid exchangeable population models. Ann Probab. 29(4):1547–1562. 10.1214/aop/1015345761

Morales-Arce AY, Johri P, Jensen JD. 2022. Inferring the distribution of fitness effects in patient-sampled and experimental virus populations: two case studies. Heredity. 128(2):79–87. 10.1038/s41437-021-00493-y

Morgan AP et al. 2020. Falciparum malaria from coastal Tanzania and Zanzibar remains highly connected despite effective control efforts on the archipelago. Malar J. 19(1):47. 10.1186/s12936-020-3137-8

Mota MM et al. 2001. Migration of *Plasmodium* sporozoites through cells before infection. Science. 291(5501):141–144. 10.1126/science.291.5501.141

Muhamad P et al. 2011. Polymorphisms of molecular markers of antimalarial drug resistance and relationship with artesunate-mefloquine combination therapy in patients with uncomplicated *Plasmodium falciparum* malaria in Thailand. Am J Trop Med Hyg. 85(3):568–72. 10.4269/ajtmh.2011.11-0194

Musiime AK et al. 2019. Is that a real oocyst? Insectary establishment and identification of *Plasmodium falciparum* oocysts in midguts of *Anopheles* mosquitoes fed on infected human blood in Tororo, Uganda. Malar J. 18(1):287. 10.1186/s12936-019-2922-8

Musto H et al. 1999. Synonymous codon choices in the extremely GC-poor genome of *Plasmodium falciparum*: compositional constraints and translational selection. J Mol Evol. 49(1):27–35. 10.1007/PL00006531

Myers S et al. 2005. A fine-scale map of recombination rates and hotspots across the human genome. Science. 310(5746):321–324. 10.1126/science.1117196

Mzilahowa T, McCall PJ, Hastings IM. 2007. “Sexual” population structure and genetics of the malaria agent *P. falciparum*. PLoS One. 2(7):e613. 10.1371/journal.pone.0000613

Nachman MW, Crowell SL. 2000. Estimate of the mutation rate per nucleotide in humans. Genetics. 156(1):297–304. 10.1093/genetics/156.1.297

Ness RW et al. 2015. Extensive de novo mutation rate variation between individuals and across the genome of *Chlamydomonas reinhardtii*. Genome Res. 25(11):1739–1749. 10.1101/gr.191494.115

Niaré K et al. 2025. A novel locus associated with decreased susceptibility of *Plasmodium falciparum* to lumefantrine and dihydroartemisinin has emerged and spread in Uganda. 10.1101/2025.07.30.667738

Nicolaisen LE, Desai MM. 2013. Distortions in genealogies due to purifying selection and recombination. Genetics. 195(1):221–230. 10.1534/genetics.113.152983

Nielsen R. 2005. Molecular signatures of natural selection. Annu Rev Genet. 39(1):197–218. 10.1146/annurev.genet.39.073003.112420

Nkhoma SC et al. 2018. Resolving within-host malaria parasite diversity using single-cell sequencing. http://biorxiv.org/lookup/doi/10.1101/391268. 10.1101/391268

Nkhoma SC et al. 2020. Co-transmission of related malaria parasite lineages shapes within-host parasite diversity. Cell Host Microbe. 27(1):93–103.e4. 10.1016/j.chom.2019.12.001

Noedl H et al. 2008. Evidence of artemisinin-resistant malaria in western Cambodia. N Engl J Med. 359(24):2619–2620. 10.1056/NEJMc0805011

Nordborg M, Charlesworth B, Charlesworth D. 1996. The effect of recombination on background selection. Genet Res. 67(2):159–174. 10.1017/s0016672300033619

Nsanzabana C et al. 2010. Quantifying the evolution and impact of antimalarial drug resistance: drug use, spread of resistance, and drug failure over a 12-year period in Papua New Guinea. J Infect Dis. 201(3):435–443. 10.1086/649784

Obaldia N et al. 2015. Clonal outbreak of *Plasmodium falciparum* infection in eastern Panama. J Infect Dis. 211(7):1087–1096. 10.1093/infdis/jiu575

Ocholla H et al. 2014. Whole-genome scans provide evidence of adaptive evolution in Malawian *Plasmodium falciparum* isolates. J Infect Dis. 210(12):1991–2000. 10.1093/infdis/jiu349

Ohta T, Kimura M. 1971. Linkage disequilibrium between two segregating nucleotide sites under the steady flux of mutations in a finite population. Genetics. 68(4):571–580. 10.1093/genetics/68.4.571

Ollivier L, Charlesworth B, Pouyet F. 2025. Beyond recombination: exploring the impact of meiotic frequency on genome-wide genetic diversity. PLoS Genet. 21(8):e1011798. 10.1371/journal.pgen.1011798

Orr HA, Otto SP. 1994. Does diploidy increase the rate of adaptation? Genetics. 136(4):1475–1480. 10.1093/genetics/136.4.1475

Osborne A et al. 2024. *Plasmodium falciparum* population dynamics in East Africa and genomic surveillance along the Kenya-Uganda border. Sci Rep. 14(1):18051. 10.1038/s41598-024-67623-4

Otto SP, Gerstein AC. 2008. The evolution of haploidy and diploidy. Curr Biol. 18(24):R1121–R1124. 10.1016/j.cub.2008.09.039

Otto TD et al. 2014. Genome sequencing of chimpanzee malaria parasites reveals possible pathways of adaptation to human hosts. Nat Commun. 5(1):4754. 10.1038/ncomms5754

Otto TD, Böhme U, et al. 2018. Long read assemblies of geographically dispersed *Plasmodium falciparum* isolates reveal highly structured subtelomeres. Wellcome Open Res. 3:52. 10.12688/wellcomeopenres.14571.1

Otto TD, Gilabert A, et al. 2018. Genomes of all known members of a *Plasmodium* subgenus reveal paths to virulent human malaria. Nat Microbiol. 3(6):687–697. 10.1038/s41564-018-0162-2

Parobek CM et al. 2016. Selective sweep suggests transcriptional regulation may underlie *Plasmodium vivax* resilience to malaria control measures in Cambodia. Proc Natl Acad Sci USA. 113(50):E8096–E8105. 10.1073/pnas.1608828113

Parobek CM et al. 2017. Partner-drug resistance and population substructuring of artemisinin-resistant *Plasmodium falciparum* in Cambodia. Genome Biol Evol. 9(6):1673–1686. 10.1093/gbe/evx126

Paschalidis A et al. 2023. coiaf: Directly estimating complexity of infection with allele frequencies. PLoS Comput Biol. 19(6):e1010247. 10.1371/journal.pcbi.1010247

Payne D. 1987. Spread of chloroquine resistance in *Plasmodium falciparum*. Parasitol Today. 3(8):241–246. 10.1016/0169-4758(87)90147-5

Peixoto L, Fernández V, Musto H. 2004. The effect of expression levels on codon usage in *Plasmodium falciparum*. Parasitology. 128(3):245–251. 10.1017/S0031182003004517

Peñalba JV, Wolf JBW. 2020. From molecules to populations: appreciating and estimating recombination rate variation. Nat Rev Genet. 21(8):476–492. 10.1038/s41576-020-0240-1

Perkins SA, Neafsey DE, Early AM. 2025. Heterogeneous constraint and adaptation across the malaria parasite life cycle. Proc R Soc B. 292(2058):20251549. 10.1098/rspb.2025.1549

Phompradit P et al. 2014. Four years’ monitoring of in vitro sensitivity and candidate molecular markers of resistance of *Plasmodium falciparum* to artesunate-mefloquine combination in the Thai-Myanmar border. Malar J. 13:23. 10.1186/1475-2875-13-23

Phyo AP et al. 2016. Declining efficacy of artemisinin combination therapy against *P. falciparum* malaria on the Thai-Myanmar border (2003-2013): The role of parasite genetic factors. Clin Infect Dis. 63(6):784–791. 10.1093/cid/ciw388

Pitman J. 1999. Coalescents with multiple collisions. Ann Probab. 27(4):1870–1902. 10.1214/aop/1022874819

Popkin-Hall ZR et al. 2024. Population genomics of *Plasmodium malariae* from four African countries. 10.1101/2024.09.07.24313132

Prieto-Baños S et al. 2025. Annotation matters: the effect of structural gene annotation on orthology inference. Bioinformatics. 41(7):btaf365. 10.1093/bioinformatics/btaf365

Prugnolle F, McGee K, Keebler J, Awadalla P. 2008. Selection shapes malaria genomes and drives divergence between pathogens infecting hominids versus rodents. BMC Evol Biol. 8(1):223. 10.1186/1471-2148-8-223

Przeworski M. 2002. The signature of positive selection at randomly chosen loci. Genetics. 160(3):1179–1189. 10.1093/genetics/160.3.1179

Ranford-Cartwright LC, Balfe P, Carter R, Walliker D. 1991. Genetic hybrids of *Plasmodium falciparum* identified by amplification of genomic DNA from single oocysts. Mol Biochem Parasitol. 49(2):239–243. 10.1016/0166-6851(91)90067-G

Razakandrainibe FG et al. 2005. “Clonal” population structure of the malaria agent *Plasmodium falciparum* in high-infection regions. Proc Natl Acad Sci USA. 102(48):17388–17393. 10.1073/pnas.0508871102

Renzette N et al. 2013. Rapid intrahost evolution of human cytomegalovirus is shaped by demography and positive selection. PLoS Genet. 9(9):e1003735. 10.1371/journal.pgen.1003735

Reuling IJ et al. 2018. A randomized feasibility trial comparing four antimalarial drug regimens to induce *Plasmodium falciparum* gametocytemia in the controlled human malaria infection model. eLife. 7:e31549. 10.7554/eLife.31549

Rogers AR, Huff C. 2009. Linkage disequilibrium between loci with unknown phase. Genetics. 182(3):839–844. 10.1534/genetics.108.093153

Rosenberg R, Rungsiwongse J. 1991. The number of sporozoites produced by individual malaria oocysts. Am J Trop Med Hyg. 45(5):574–577. 10.4269/ajtmh.1991.45.574

Rosenthal PJ et al. 2024. The emergence of artemisinin partial resistance in Africa: how do we respond? Lancet Infect Dis. 24(9):e591–e600. 10.1016/S1473-3099(24)00141-5

Roze D, Rousset F, Michalakis Y. 2005. Germline bottlenecks, biparental inheritance and selection on mitochondrial variants. Genetics. 170(3):1385–1399. 10.1534/genetics.104.039495

Ruderfer DM, Pratt SC, Seidel HS, Kruglyak L. 2006. Population genomic analysis of outcrossing and recombination in yeast. Nat Genet. 38(9):1077–81. 10.1038/ng1859

Rutledge GG et al. 2017. *Plasmodium malariae* and *P. ovale* genomes provide insights into malaria parasite evolution. Nature. 542(7639):101–104. 10.1038/nature21038

Sabin S, Morales-Arce AY, Pfeifer SP, Jensen JD. 2022. The impact of frequently neglected model violations on bacterial recombination rate estimation: a case study in *Mycobacterium canettii* and *Mycobacterium tuberculosis*. G3. 12(5):jkac055. 10.1093/g3journal/jkac055

Sáenz FE et al. 2015. Clonal population expansion in an outbreak of *Plasmodium falciparum* on the northwest coast of Ecuador. Malar J. 14(1):497. 10.1186/s12936-015-1019-2

Samad H et al. 2015. Imputation-based population genetics analysis of *Plasmodium falciparum* malaria parasites. PLoS Genet. 11(4):e1005131. 10.1371/journal.pgen.1005131

Santiago E, Caballero A. 1998. Effective size and polymorphism of linked neutral loci in populations under directional selection. Genetics. 149(4):2105–2117. 10.1093/genetics/149.4.2105

dos Santos G, et al. 2015. FlyBase: introduction of the *Drosophila melanogaster* release 6 reference genome assembly and large-scale migration of genome annotations. Nucleic Acids Res. 43(D1):D690–D697. 10.1093/nar/gku1099

Sato S. 2021. *Plasmodium* —a brief introduction to the parasites causing human malaria and their basic biology. J Physiol Anthropol. 40:1. 10.1186/s40101-020-00251-9

Saunders DL, Vanachayangkul P, Lon C. 2014. Dihydroartemisinin-piperaquine failure in Cambodia. N Engl J Med. 371(5):484–485. 10.1056/NEJMc1403007

Scott TW, Takken W. 2012. Feeding strategies of anthropophilic mosquitoes result in increased risk of pathogen transmission. Trends Parasitol. 28(3):114–121. 10.1016/j.pt.2012.01.001

Segovia X et al. 2025. Assessing fitness costs in malaria parasites: a comprehensive review and implications for drug resistance management. Malar J. 24(1):65. 10.1186/s12936-025-05286-w

Sheehan S, Song YS. 2016. Deep learning for population genetic inference. PLoS Comput Biol. 12(3):e1004845. 10.1371/journal.pcbi.1004845

Shiao S-H et al. 2006. *Fz2* and *cdc42* mediate melanization and actin polymerization but are dispensable for *Plasmodium* killing in the mosquito midgut. PLoS Pathog. 2(12):e133. 10.1371/journal.ppat.0020133

Siepel A et al. 2005. Evolutionarily conserved elements in vertebrate, insect, worm, and yeast genomes. Genome Res. 15(8):1034–1050. 10.1101/gr.3715005

Silvestrini F, Alano P, Williams JL. 2000. Commitment to the production of male and female gametocytes in the human malaria parasite *Plasmodium falciparum*. Parasitology. 121(5):465–471. 10.1017/s0031182099006691

Sinha I, Woodrow CJ. 2018. Forces acting on codon bias in malaria parasites. Sci Rep. 8(1):15984. 10.1038/s41598-018-34404-9

Slatkin M. 1977. Gene flow and genetic drift in a species subject to frequent local extinctions. Theor Popul Biol. 12(3):253–262. 10.1016/0040-5809(77)90045-4

Smith LM et al. 2020. An intrinsic oscillator drives the blood stage cycle of the malaria parasite, *Plasmodium falciparum*. Science. 368(6492):754. 10.1126/science.aba4357

Smith RC, Jacobs-Lorena M. 2010. *Plasmodium*–mosquito interactions: a tale of roadblocks and detours. Adv In Insect Phys. 39:119–149. 10.1016/B978-0-12-381387-9.00004-X

Smith RC, Vega-Rodríguez J, Jacobs-Lorena M. 2014. The *Plasmodium* bottleneck: malaria parasite losses in the mosquito vector. Mem Inst Oswaldo Cruz. 109(5):644–661. 10.1590/0074-0276130597

Smith TCA, Arndt PF, Eyre-Walker A. 2018. Large scale variation in the rate of germ-line de novo mutation, base composition, divergence and diversity in humans. PLoS Genet. 14(3):e1007254. 10.1371/journal.pgen.1007254

Smith TG et al. 2000. Commitment to sexual differentiation in the human malaria parasite, *Plasmodium falciparum*. Parasitology. 121(2):127–133. 10.1017/s0031182099006265

Snounou G, Beck H-P. 1998. The use of PCR genotyping in the assessment of recrudescence or reinfection after antimalarial drug treatment. Parasitol Today. 14(11):462–467. 10.1016/S0169-4758(98)01340-4

Snow RW. 2015. Global malaria eradication and the importance of *Plasmodium falciparum* epidemiology in Africa. BMC Med. 13:23. 10.1186/s12916-014-0254-7

Soni V, Jensen JD. 2024. Temporal challenges in detecting balancing selection from population genomic data. G3. 14(6):jkae069. 10.1093/g3journal/jkae069

Soni V, Johri P, Jensen JD. 2023. Evaluating power to detect recurrent selective sweeps under increasingly realistic evolutionary null models. Evolution. 77(10):2113–2127. 10.1093/evolut/qpad120

Städler T et al. 2009. The impact of sampling schemes on the site frequency spectrum in nonequilibrium subdivided populations. Genetics. 182(1):205–216. 10.1534/genetics.108.094904

Stephan W. 2019. Selective sweeps. Genetics. 211(1):5–13. 10.1534/genetics.118.301319

Stokes BH et al. 2021. *Plasmodium falciparum* K13 mutations in Africa and Asia impact artemisinin resistance and parasite fitness. eLife. 10:e66277. 10.7554/eLife.66277

Su X et al. 1999. A genetic map and recombination parameters of the human malaria parasite *Plasmodium falciparum*. Science. 286(5443):1351–1353. 10.1126/science.286.5443.1351

Tadesse FG et al. 2018. The relative contribution of symptomatic and asymptomatic *Plasmodium vivax* and *Plasmodium falciparum* infections to the infectious reservoir in a low-endemic setting in Ethiopia. Clinical Infectious Diseases. 66(12):1883–1891

Tadesse FG et al. 2019. Gametocyte sex ratio: the key to understanding *Plasmodium falciparum* transmission? Trends Parasitol. 35(3):226–238. 10.1016/j.pt.2018.12.001

Tajima F. 1989. Statistical method for testing the neutral mutation hypothesis by DNA polymorphism. Genetics. 123(3):585–595. 10.1093/genetics/123.3.585

Takala-Harrison S, Laufer MK. 2015. Antimalarial drug resistance in Africa: key lessons for the future. Ann NY Acad Sci. 1342:62–67. 10.1111/nyas.12766

Taylor AR et al. 2017. Quantifying connectivity between local *Plasmodium falciparum* malaria parasite populations using identity by descent. PLOS Genetics. 13(10):e1007065. 10.1371/journal.pgen.1007065

Tellier A, Lemaire C. 2014. Coalescence 2.0: a multiple branching of recent theoretical developments and their applications. Mol Ecol. 23(11):2637–2652. 10.1111/mec.12755

Teshima KM, Coop G, Przeworski M. 2006. How reliable are empirical genomic scans for selective sweeps? Genome Res. 16(6):702–712. 10.1101/gr.5105206

Thanh NV et al. 2017. Rapid decline in the susceptibility of *Plasmodium falciparum* to dihydroartemisinin-piperaquine in the south of Vietnam. Malar J. 16(1):27. 10.1186/s12936-017-1680-8

Thornton KR, Jensen JD. 2007. Controlling the false-positive rate in multilocus genome scans for selection. Genetics. 175(2):737–750. 10.1534/genetics.106.064642

Triglia T, Menting JGT, Wilson C, Cowman AF. 1997. Mutations in dihydropteroate synthase are responsible for sulfone and sulfonamide resistance in *Plasmodium falciparum*. Proc Natl Acad Sci USA. 94(25):13944–13949. 10.1073/pnas.94.25.13944

Uyoga S et al. 2021. Plasma *Plasmodium falciparum* histidine-rich protein 2 concentrations in children with malaria infections of differing severity in Kilifi, Kenya. Clin Infect Dis. 73(7):e2415–e2423. 10.1093/cid/ciaa1141

Vaughan AM et al. 2012. Complete *Plasmodium falciparum* liver-stage development in liver-chimeric mice. J Clin Invest. 122(10):3618–3628. 10.1172/JCI62684

Vaughan AM, Kappe SHI. 2017. Malaria parasite liver infection and exoerythrocytic biology. Cold Spring Harb Perspect Med. 7(6):a025486. 10.1101/cshperspect.a025486

Venugopal K, Hentzschel F, Valkiūnas G, Marti M. 2020. *Plasmodium* asexual growth and sexual development in the haematopoietic niche of the host. Nat Rev Microbiol. 18(3):177–189. 10.1038/s41579-019-0306-2

Viriyakosol S et al. 1995. Genotyping of *Plasmodium falciparum* isolates by the polymerase chain reaction and potential uses in epidemiological studies. Bull World Health Organ. 73(1):85–95

Volkman SK et al. 2007. A genome-wide map of diversity in *Plasmodium falciparum*. Nat Genet. 39(1):113–119. 10.1038/ng1930

Wakeley J, Aliacar N. 2001. Gene genealogies in a metapopulation. Genetics. 159(2):893–905. 10.1093/genetics/159.2.893

Walker-Jonah A et al. 1992. An RFLP map of the *Plasmodium falciparum* genome, recombination rates and favored linkage groups in a genetic cross. Mol Biochem Parasitol. 51(2):313–320. 10.1016/0166-6851(92)90081-T

Walliker D et al. 1987. Genetic analysis of the human malaria parasite *Plasmodium falciparum*. Science. 236(4809):1661–1666. 10.1126/science.3299700

Walliker D, Hunt P, Babiker H. 2005. Fitness of drug-resistant malaria parasites. Acta Trop. 94(3):251–259. 10.1016/j.actatropica.2005.04.005

Wang CYT et al. 2018. Assessing *Plasmodium falciparum* transmission in mosquito-feeding assays using quantitative PCR. Malar J. 17(1):249. 10.1186/s12936-018-2382-6

Wang Z et al. 2016. Genome-wide association analysis identifies genetic loci associated with resistance to multiple antimalarials in *Plasmodium falciparum* from China-Myanmar border. Sci Rep. 6(1):33891. 10.1038/srep33891

Watson OJ et al. 2017. Modelling the drivers of the spread of *Plasmodium falciparum hrp2* gene deletions in sub-Saharan Africa. eLife. 6:e25008. 10.7554/eLife.25008

Weedall GD, Hall N. 2015. Sexual reproduction and genetic exchange in parasitic protists. Parasitology. 142 Suppl 1(Suppl 1):S120–S127. 10.1017/S0031182014001693

Wellems TE, Plowe CV. 2001. Chloroquine-resistant malaria. J Infect Dis. 184(6):770–776. 10.1086/322858

White NJ. 2017. Malaria parasite clearance. Malar J. 16(1):88. 10.1186/s12936-017-1731-1

Whitlock AOB, Juliano JJ, Mideo N. 2021. Immune selection suppresses the emergence of drug resistance in malaria parasites but facilitates its spread. PLoS Comput Biol. 17(7):e1008577. 10.1371/journal.pcbi.1008577

Wootton JC. 1994. Non-globular domains in protein sequences: automated segmentation using complexity measures. Comput Chem. 18(3):269–285. 10.1016/0097-8485(94)85023-2

Wootton JC et al. 2002. Genetic diversity and chloroquine selective sweeps in *Plasmodium falciparum*. Nature. 418(6895):320–323. 10.1038/nature00813

World Health Organization. 2024. [accessed 2025 Aug 19]. https://www.who.int/teams/global-malaria-programme/reports/world-malaria-report-2024

Wright S. 1943. Isolation by distance. Genetics. 28(2):114–138. 10.1093/genetics/28.2.114

Xu P et al. 2026. e3SIM: Epidemiological-ecological-evolutionary simulation framework for genomic epidemiology. Methods Ecol Evol. 17(4):1051–1059. 10.1111/2041-210x.70278

Yamauchi LM, Coppi A, Snounou G, Sinnis P. 2007. *Plasmodium* sporozoites trickle out of the injection site. Cell Microbiol. 9(5):1215–1222. 10.1111/j.1462-5822.2006.00861.x

Zhang M et al. 2018. Uncovering the essential genes of the human malaria parasite *Plasmodium falciparum* by saturation mutagenesis. Science. 360(6388):eaap7847. 10.1126/science.aap7847

Zilversmit MM et al. 2010. Low-complexity regions in *Plasmodium falciparum*: missing links in the evolution of an extreme genome. Mol Biol Evol. 27(9):2198–2209. 10.1093/molbev/msq108

