## Supplementary Information for "Towards an evolutionary baseline model of *Plasmodium falciparum* for population-genomic inference"

### SUPPLEMENTARY FIGURES AND TABLES

**Table S1**: Previous estimates of the duration of parts of the lifecycle of *P. falciparum*, their approximate population sizes, and ploidy during the various stages of their life cycle. Variation in these estimates is indicated by the column "Overall range in the literature".

| **Stage** | **Published estimates** | **Overall range in the literature** | **Length of time** | **Ploidy** |
|---|---|---|---|---|
| Dermis (Human) | **$10^2$** (order-of-magnitude ( review synthesis); Graumans et al. 2020); **116** (geometric mean; IQR: 33-501; Andolina et al. 2024) **432** (geometric mean; IQR:116-2,779; Andolina et al. 2024) **862** (mean; range: 354-2,666; (Kanatani et al. 2024) | $10^2$-$10^3$ | 1-3hr | *N* (Mzilahowa et al. 2007) |
| Pre-hepatocyte Erythrocytic stage (Human) | **10-$10^2$** (order-of-magnitude (review synthesis); Kappe et al. 2010) **1-10** (order-of-magnitude (review synthesis); Graumans et al. 2020) | 1-$10^2$ | 60s (Cowman and Crabb 2006) | *N* (Mzilahowa et al. 2007) |
| Liver (Human) | **$10^4$** (order-of-magnitude (review synthesis); Graumans et al. 2020) **$10^4$-$10^5$** (order-of-magnitude (review synthesis); Kappe et al. 2010) | $10^4 - 10^5$ | 6-7 days (Vaughan et al. 2012) | *N* (Mzilahowa et al. 2007) |

| | | | | |
|---|---|---|---|---|
| | **90,000** (max estimate; (Vaughan et al. 2012; Vaughan and Kappe 2017) | | | |
| Asexual blood stage (Human)* | $\mathbf{10^{11}}$ (order-of-magnitude (review synthesis); Graumans et al. 2020)<br>$\mathbf{10^{9}}$-$\mathbf{10^{13}}$ (order-of-magnitude (review synthesis); Kappe et al. 2010)<br>$\mathbf{3.6 \times 10^{10}}$ * (symptomatic median; IQR: $3.4\times10^{9}$-$6.4\times10^{11}$; Tadesse et al. 2018)<br>$\mathbf{10^{8}}$ * (asymptomatic median; IQR: $3\times10^{6}$-$2\times10^{9}$; Tadesse et al. 2018) | $10^{6}-10^{13}$ | 48-hr cycles (Hawking et al. 1968; Smith et al. 2020) | *N* (Mzilahowa et al. 2007) |
| Gametocytes (Human) | $\mathbf{10^{10}}$ (order-of-magnitude (review synthesis); Graumans et al. 2020)<br>$\mathbf{10^{6}-10^{9}}$ (inferred order-of-magnitude; review synthesis; (Lin et al. 2014) | $10^{6}-10^{10}$ | 9-12 days (Hawking et al. 1968; Venugopal et al. 2020) | *N* (Mzilahowa et al. 2007) |

| | | | | |
|---|---|---|---|---|
| Gametocytes (Mosquito) | **$10\text{-}10^3$** (order-of-magnitude (review synthesis); Kappe et al. 2010)<br>**443.5** (mean; range: 64-2392; Gouagna et al. 1998; reviewed in Smith et al. 2014)<br>$\mathbf{1-10^3}$ * (range (inferred quantity); Lin et al. 2014) | $1\text{-}10^3$ | ~15 mins for maturity (Sinden et al. 1978)<br>~15 mins (Smith et al. 2014) | $N$ (Mzilahowa et al. 2007) |
| Gametes (Mosquito) | **$10^3$** (order-of-magnitude (review synthesis); Graumans et al. 2020) | $10^3$ | ~45mins for fusion (Smith et al. 2014) | $N$ (Mzilahowa et al. 2007) |
| Zygote (Mosquito) | **$10^2$** (order-of-magnitude (review synthesis); Graumans et al. 2020)<br>**12.6** (mean; range: 0.2-77.3; Gouagna et al. 1998; reviewed in Smith et al. 2014) | $10\text{-}10^2$ | 17-23hr (Smith et al. 2014) | $2N$ (Graumans et al. 2020) |
| Ookinite (Mosquito) | **$10^2$** (order-of-magnitude (review synthesis); Graumans et al. 2020)<br>**5.5** (mean; range: 0-35.7; Gouagna et al. 1998; reviewed in Smith et al. 2014) | $10\text{-}10^2$ | 17-23hr | $4N$ (Mzilahowa et al. 2007) |

| | | | | |
|---|---|---|---|---|
| Oocyst (Mosquito) | **1-10** (Kappe et al. 2010)<br>**1-10** (Graumans et al. 2020)<br>**2** (mean; range: 0-16.5; Gouagna et al. 1998; reviewed in Smith et al. 2014)<br>**5-6** (median; IQR: 2-20; (Andolina et al. 2014) | 1-10 | ~9-14 days for development + spore production (Smith et al. 2014) | *N* (Mzilahowa et al. 2007) |
| Sporozoites per oocyst (Mosquito) | **3,600** (median; IQR:2720-5453; (Wang et al. 2018)<br>**3,385** (mean; range:359-4,554) (Rosenberg and Rungsiwongse 1991)<br>**9604** (median; range: 2489-14,520) (Kanatani et al. 2024)<br>**15,390** (IQR: 10,600-20,887) (Andolina et al. 2024)<br>**1500-5000** (Graumans et al. 2020) | $10^3 - 10^4$ | ~9-14 days for development + spore production (Smith et al. 2014) | *N* (Mzilahowa et al. 2007) |
| Salivary gland (Mosquito) | **21,016** (mean (lab); IQR: 9137-78,380; (Andolina et al. 2024)<br>**35,149** (mean (field); IQR: 20,310-164,900; Andolina et al. 2024)<br>**$10^3$-$10^4$** (Kappe et al. 2010)<br>**$10^4$** (Graumans et al. 2020)<br>**~$10^4$** **(derived median range; range: $3 \times 10^3 - 1 \times 10^5$; Kanatani et al. 2024) | $10^3$-$10^5$ | 2-4 days (Scott and Takken 2012; Brackney et al. 2021) | *N* (Mzilahowa et al. 2007) |

*Total body individuals were estimated by taking the reported density (parasites/microliter) and multiplying by 5L of blood.

**To estimate the salivary gland sporozoite population, the reported median fecundity of 4,236 salivary gland sporozoites per ruptured oocyst (IQR: 2,997–11,542) were used. Then a general representative population size was estimated by multiplying this median by the typical number of ruptured oocysts (3–6). To derive a plausible range based on the number of oocysts, the lower quartile was combined with a single ruptured oocyst (2,997 × 1 oocyst ~~$3 \times 10^3$) and the upper quartile with 10 ruptured oocysts (11,542 × 10 ~ $1.15 \times 10^5$). The salivary gland sporozoite burdens were found to be highly right-skewed. So, this upper bound would cover the extreme tail of a highly infected mosquito.

**Table S2**: The proportion of time spent in each ploidy.

| Stage | Ploidy | Minimum Time (hr) | Maximum Time (hr) |
|---|---|---|---|
| Dermis (skin) | *N* | 1 | 3 |
| Pre-liver | *N* | 0.0167 | 0.0167 |
| Liver | *N* | 144 | 168 |
| Gametocyte | *N* | 216 | 288 |
| Gamete | *N* | 0.25 | 0.25 |
| Gamete Fusion | 2*N* | 0.75 | 0.75 |
| Ookinite | 4*N* | 17 | 23 |
| Oocyst | *N* | 216 | 336 |
| Salivary Gland | *N* | 48 | 96 |

| | *N* Minimum (hr) | *N* Maximum (hr) | 2*N* Minimum (hr) | 2*N* Maximum (hr) | 4*N* Minimum (hr) | 4*N* Maximum (hr) |
|---|---|---|---|---|---|---|
| Sum | 625.27 | 891.27 | 0.75 | 0.75 | 17 | 23 |
| Percentage | 97.24% | 97.40% | 0.12% | 0.082% | 2.64% | 2.51% |

**Table S3.** Mean observed *P. falciparum Fws* values for all countries in the MalariaGEN Pf7 (2023)

| Country | Number of Samples | Mean Fws |
|---|---|---|
| Bangladesh | 1310 | 0.91 |
| Benin | 150 | 0.895 |
| Burkina Faso | 57 | 0.733 |
| Cambodia | 1267 | 0.948 |
| Cameroon | 264 | 0.825 |
| Colombia | 135 | 0.979 |
| Côte d'Ivoire | 71 | 0.882 |
| Democratic Republic of the Congo | 520 | 0.817 |
| Ethiopia | 21 | 0.99 |
| Gabon | 55 | 0.886 |
| Gambia | 863 | 0.879 |
| Ghana | 3131 | 0.825 |
| Guinea | 151 | 0.812 |
| India | 300 | 0.896 |
| Indonesia | 121 | 0.943 |
| Kenya | 690 | 0.838 |
| Laos | 991 | 0.964 |
| Madagascar | 24 | 0.954 |
| Malawi | 265 | 0.751 |
| Mali | 1167 | 0.861 |
| Mauritania | 92 | 0.878 |
| Mozambique | 34 | 0.867 |
| Myanmar | 985 | 0.978 |
| Nigeria | 110 | 0.923 |
| Papua New Guinea | 221 | 0.975 |
| Peru | 21 | 0.999 |
| Senegal | 150 | 0.942 |
| Sudan | 76 | 0.966 |

| | | |
|---|---|---|
| Tanzania | 589 | 0.851 |
| Thailand | 954 | 0.941 |
| Uganda | 12 | 0.814 |
| Venezuela | 2 | 0.892 |
| Vietnam | 1404 | 0.962 |

**Table S4**: Theoretical expectations of neutral nucleotide diversity with BGS. Here we assume a Wright-Fisher population size of 80,000 haploid individuals and a mutation rate of $5.6 \times 10^{-9}$ per site/ generation. We assumed $R_{eff} = R(1 - F_{it})$ where $F = 0.6$ for high-transmission and 0.9 for low-transmission populations, where $F$ is the inbreeding coefficient. The genome-wide deleterious mutation rate ($U$) is obtained by assuming that 53% of the genome is under selection. When assuming a single selection coefficient, the relative diversity with BGS to that under neutrality, depicted by $B$, was calculated as $B \sim exp\{-U/(2s + R_{self}(1 - s))\}$. When assuming that selection coefficients follow a distribution of fitness effects (DFE), a combination of four non-overlapping uniform distributions was assumed, such that $0<Ns\leq1$, $1<Ns\leq10$, $10<Ns\leq100$, and $100 <Ns\leq100{,}000$, where $s$ represents the disadvantage of a homozygous mutant relative to wild type. Each non-overlapping distribution was present in proportions $f_0$, $f_1$, $f_2$, and $f_3$, respectively. We assumed that $f_0$=0, $f_1$=0.333, $f_2$=0.333, $f_3$=0.333 at selected sites. When assuming a DFE, $B$ was estimated using equations described in Marsh et al. (2025).
.

| | **Smallest chromosome** | **Average chromosome** | **Longest chromosome** |
|---|---|---|---|
| Length | 640 kb | 1.6 Mb | 3.3 Mb |
| $U$ (from base subs) | 0.002 | 0.005 | 0.010 |
| $U$ (from indels) | 0.001 | 0.002 | 0.004 |
| $U$ (total) | 0.003 | 0.007 | 0.014 |
| $R$ (in Morgans) | 0.5 | 1 | 2.25 |
| *In high-transmission populations* ($F$=0.6) | | | |
| $R_{self}$ | 0.2 | 0.4 | 0.9 |
| $B$ with strong selection ($s$=-0.1) | 0.992 | 0.989 | 0.987 |
| $B$ with weak selection ($s$=-0.01) | 0.986 | 0.986 | 0.986 |
| $B$ from linked regions (with a DFE) | 0.994 | 0.992 | 0.992 |
| $B$ from unlinked chromosomes | 0.961 | 0.963 | 0.966 |
| $B$ from linked and unlinked effects | 0.955 | 0.955 | 0.958 |
| *In low transmission populations* ($F$=0.9) | | | |
| $R_{self}$ | 0.05 | 0.1 | 0.225 |
| $B$ with strong selection ($s$=-0.1) | 0.988 | 0.980 | 0.968 |

| | | | |
|---|---|---|---|
| $B$ with weak selection ($s$=-0.01) | 0.958 | 0.951 | 0.950 |
| $B$ from linked regions (with a DFE) | 0.968 | 0.966 | 0.963 |
| $B$ from unlinked chromosomes | 0.961 | 0.963 | 0.966 |
| $B$ from linked and unlinked effects | 0.930 | 0.930 | 0.930 |

### SUPPLEMENTARY NOTE A

**Estimation of the lower and upper bounds of the number of mitotic divisions during the human blood phase**

To estimate the lower bound, we estimated the number of mitotic divisions in the human blood that would be needed to reach $10^9$ $body\ individuals$ from 100,000 number of parasites in the liver, and found that to be approximately 14 divisions. 14 divisions correspond to approximately 3 48-hour cycles, or 6 days. This estimation is concordant with the estimated number of days to the blood density threshold for symptom onset in White (2017). To estimate the upper bound, we assumed ~8 days to the onset of symptoms plus ~7 days for full parasite clearance with treatment, assuming 5 divisions per 48-hour cycle(White 2017; McDew-White et al. 2019). This resulted in a total of 40 divisions.

### REFERENCES


Abdel Hamid MM, Abdelraheem MH, Acheampong DO, Ahouidi A, Ali M, Almagro-Garcia J, Amambua-Ngwa A, Amaratunga C, Amenga-Etego L, Andagalu B, et al. 2023. Pf7: an open dataset of *Plasmodium falciparum* genome variation in 20,000 worldwide samples [version 1; peer review: 3 approved]. *Wellcome Open Res.* 8:22.

Andolina C, Graumans W, Guelbeogo M, Gemert GJ van, Ramjith J, Harouna S, Soumanaba Z, Stoter R, Vegte-Bolmer M, Pangos M, et al. 2024. A transmission bottleneck for malaria? Quantification of sporozoite expelling by *Anopheles* mosquitoes infected with laboratory and naturally circulating *P. falciparum* gametocytes. *eLife* 12:RP90989.

Brackney DE, LaReau JC, Smith RC. 2021. Frequency matters: How successive feeding episodes by blood-feeding insect vectors influences disease transmission. *PLoS Pathog.* 17:e1009590.

Cowman AF, Crabb BS. 2006. Invasion of red blood cells by malaria parasites. *Cell* 124:755–766.

Gouagna LC, Gouagna LC, Mulder B, Mulder B, Noubissi E, Noubissi E, Tchuinkam T, Tchuinkam T, and Boudin C, Boudin C, et al. 1998. The early sporogonic cycle of *Plasmodium falciparum* in laboratory-infected *Anopheles gambiae*: an estimation of parasite efficacy. *Trop Med Int Health* 3:21–28.

Graumans W, Jacobs E, Bousema T, Sinnis P. 2020. When is a *Plasmodium*-infected mosquito an infectious mosquito? *Trends Parasitol.* 36:705–716.

Hawking F, Worms MJ, Gammage K. 1968. 24- and 48-hour cycles of malaria parasites in the blood; their purpose, production and control. *Trans. R. Soc. Trop. Med. Hyg.* 62:731–760.

Kanatani S, Stiffler D, Bousema T, Yenokyan G, Sinnis P. 2024. Revisiting the Plasmodium sporozoite inoculum and elucidating the efficiency with which malaria parasites progress through the mosquito. *Nat Commun* 15:748.

Kappe SHI, Vaughan AM, Boddey JA, Cowman AF. 2010. That was then but this is now: Malaria research in the time of an eradication agenda. *Science* 328:862–866.

Lin JT, Saunders DL, Meshnick SR. 2014. The role of submicroscopic parasitemia in malaria transmission: what is the evidence? *Trends Parasitol.* 30:183–190.

Marsh JI, Kaushik S, Johri P. 2025. Effects of rescaling forward-in-time population genetic simulations. :2025.04.24.650500. Available from: https://www.biorxiv.org/content/10.1101/2025.04.24.650500v1

McDew-White M, Li X, Nkhoma SC, Nair S, Cheeseman I, Anderson TJC. 2019. Mode and tempo of microsatellite length change in a malaria parasite mutation accumulation experiment. *Genome Biol. Evol.* 11:1971–1985.

Mzilahowa T, McCall PJ, Hastings IM. 2007. “Sexual” population structure and genetics of the malaria agent *P. falciparum*. *PLoS One* 2:e613.

Rosenberg R, Rungsiwongse J. 1991. The number of sporozoites produced by individual malaria oocysts. *Am. J. Trop. Med. Hyg.* 45:574–577.

Scott TW, Takken W. 2012. Feeding strategies of anthropophilic mosquitoes result in increased risk of pathogen transmission. *Trends Parasitol.* 28:114–121.

Sinden RE, Canning EU, Bray RS, Smalley ME. 1978. Gametocyte and gamete development in *Plasmodium falciparum*. *Proc. R. Soc. Lond.* 201:375–399.

Smith LM, Motta F, Chopra G, Moch JK, Nerem RR, Cummins B, Roche KE, Kelliher CM, Leman AR, Harer J, et al. 2020. An intrinsic oscillator drives the blood stage cycle of the malaria parasite, *Plasmodium falciparum*. *Science* 368:754.

Smith RC, Vega-Rodríguez J, Jacobs-Lorena M. 2014. The *Plasmodium* bottleneck: malaria parasite losses in the mosquito vector. *Mem. Inst. Oswaldo Cruz* 109:644–661.

Tadesse FG, Slater HC, Chali W, Teelen K, Lanke K, Belachew M, Menberu T, Shumie G, Shitaye G, Okell LC, et al. 2018. The relative contribution of symptomatic and asymptomatic *Plasmodium vivax* and *Plasmodium falciparum* infections to the infectious reservoir in a low-endemic setting in Ethiopia. *Clinical Infectious Diseases* 66:1883–1891.

Vaughan AM, Kappe SHI. 2017. Malaria parasite liver infection and exoerythrocytic biology. *Cold Spring Harb. Perspect. Med.* 7:a025486.

Vaughan AM, Mikolajczak SA, Wilson EM, Grompe M, Kaushansky A, Camargo N, Bial J, Ploss A, Kappe SHI. 2012. Complete *Plasmodium falciparum* liver-stage development in liver-chimeric mice. *J. Clin. Invest.* 122:3618–3628.

Venugopal K, Hentzschel F, Valkiūnas G, Marti M. 2020. *Plasmodium* asexual growth and sexual development in the haematopoietic niche of the host. *Nat. Rev. Microbiol.* 18:177–189.

Wang CYT, McCarthy JS, Stone WJ, Bousema T, Collins KA, Bialasiewicz S. 2018. Assessing *Plasmodium falciparum* transmission in mosquito-feeding assays using quantitative PCR. *Malar. J.* 17:249.

White NJ. 2017. Malaria parasite clearance. *Malar J* 16:88.
